# Kin-dependent dispersal influences relatedness and genetic structuring in a lek system

**DOI:** 10.1101/518829

**Authors:** Hugo Cayuela, Laurent Boualit, Martin Laporte, Jérôme G. Prunier, Françoise Preiss, Alain Laurent, Francesco Foletti, Jean Clobert, Gwenaël Jacob

**Author notes:** Equal contribution (Hugo Cayuela, Laurent Boualit). GJ, JC and HC conceived the ideas and designed methodology; FP and AL collected the data; HC, ML, FF, JGP and LB analysed the data; HC led the writing of the manuscript. All authors contributed critically to the drafts and gave final approval for publication.

## Abstract

Kin selection and dispersal play a critical role in the evolution of cooperative breeding systems. Limited dispersal dramatically increases relatedness in spatially structured populations (population viscosity), with the result that neighbours tend to be genealogical relatives. Yet the increase in neighbours’ performance through altruistic interaction may also result in habitat saturation and thus exacerbate local competition between kin. Our goal was to detect the footprint of kin selection and competition by examining the spatial structure of relatedness and by comparing non-effective and effective dispersal in a population of a lekking bird, *Tetrao urogallus*. For this purpose, we analysed capture–recapture and genetic data collected over a 6-year period on a spatially structured population of *T. urogallus* in France. Our findings revealed a strong spatial structure of relatedness in males. They also indicated that the population viscosity allowed male cooperation through two non-exclusive mechanisms. First, at their first lek attendance, males aggregate in a lek composed of relatives. Second, the distance corresponding to non-effective dispersal dramatically outweighed effective dispersal distance, which suggests that dispersers incur high post-settlement costs. These two mechanisms result in strong population genetic structuring in males. In females, our findings revealed a lower level of spatial structure of relatedness and genetic structure in respect to males. Additionally, non-effective dispersal and effective dispersal distances in females were highly similar, which suggests limited post-settlement costs. These results indicate that kin-dependent dispersal decisions and costs are factors driving the evolution of cooperative courtship and have a genetic footprint in wild populations.

## Introduction

Kin selection plays a critical role in the evolution of cooperative breeding systems (Clutton-Brock 2002, Griffin & West 2003, Bourke 2011). In these systems, breeding individuals are assisted by cooperative individuals (also called helpers) that apparently sacrifice part or all of their own reproductive potential. An explanation for this apparent paradox is put forward by Hamilton’s theory (1964): because relatives share genes in common, if an individual improves the reproductive success of its relatives, this may increase its overall genetic contribution to future generations (i.e. inclusive fitness). In turn, genes that promote cooperation may then increase in frequency in the population. Accordingly, relatedness seems a prerequisite for the evolution of cooperative behaviour, implying that individuals should tend to interact within family groups or with other genetically similar individuals. Two alternative mechanisms have been proposed regarding how an individual ensures that cooperative behaviour is directed primarily towards social partners with high relatedness. The first is the ability to recognize and act preferentially towards genealogical kin (kin discrimination), which requires odour-based recognition systems or sophisticated cognitive faculties (Hepper 2005, Komdeur et al. 2008). The second is limited dispersal, which drastically increases relatedness in spatially structured populations (population viscosity) (Hamilton 1964, El Mouden & Gardner 2008, Platt & Bever 2009), with the result that neighbours tend to be genealogical relatives. Yet the increase in neighbours’ performance through altruistic interaction may also result in habitat saturation and thus exacerbate local competition between kin (Queller 1992, Platt & Bever 2009). In general, the evolution of sociality is based on a balance between kin cooperation and kin competition, which depends on a species’ life history traits, the spatial scale over which cooperation and competition occur, and the underlying habitat structure (Queller 1992; Le Galliard et al. 2003, 2004).

Dispersal designates the movement of an individual from its birth site to its reproduction patch, as well as between successive reproduction sites (Clobert et al. 2009, Matthysen 2012). It is usually considered a three-stage process that includes departure (or emigration), transience (or transfer within the landscape matrix) and settlement (or immigration) (Ronce 2007, Baguette et al. 2013). A recurrent finding of theoretical models is that dispersal evolution is based on a trade-off between costs (i.e. energetic costs, time costs, risk costs and opportunity costs) and benefits at each stage of the process (Bonte et al. 2012). In cooperative breeding systems, relatively high dispersal rates have been reported by some field studies (e.g. Greenwood & Harvey 1982, Clarke et al. 1997), a phenomenon that should decrease the relatedness level within a patch, reduce the viscosity of the whole population and constrain the evolution of cooperative behaviour. Two explanations may be possible for this apparent paradox: first, it is conceivable that population viscosity is maintained if dispersers pay acute dispersal costs after settling into a new patch (Forero et al. 2002, Hansson et al. 2004, Nystrand 2007, Dickinson et al. 2009), limiting their contribution to local reproduction. Second, cooperative behaviour may evolve if dispersers preferentially join patches composed of close relatives (Sinervo & Clobert 2003). To date, little is known about the consequences of dispersal costs and kin-dependent dispersal on gene flow and population viscosity, and therefore about their contribution to the evolution of cooperative behaviour.

Lek mating systems are an interesting study case for investigating this issue. In lek-breeding species, males congregate on display grounds during the breeding period, allowing females to compare potential partners before soliciting mating. A female’s fitness is mainly limited by her own reproductive investment. Females are thus under selection pressure to select high-quality partners, which in turn imposes a sexual selection pressure on males to advertise their quality through costly ornamental traits and courtship (Møller & Alatalo 1999, Tregenza & Wedell 2000). This leads to intense intrasexual competition and strongly skewed reproductive success in males (Höglund & Alatalo 2014, Verkuil et al. 2014). In several lek-breeding birds, theoretical studies have demonstrated that low-quality males might benefit from cooperative courtship through the indirect benefits of kin selection (Kokko & Lindstrom 1996). Empirical studies have shown that females preferentially mate in large leks (Alatalo et al. 1992). By joining a lek occupied by relatives, males with low reproduction prospects may enhance the overall attractiveness of the lek and therefore increase the fitness of related dominant males (Höglund 2003). The males unsuccessful in obtaining copulations may themselves benefit from the resulting inclusive fitness (Krakauer 2005, DuVal 2007). Theoretically, limited dispersal between leks should ensure a sufficient level of population viscosity to favour the evolution of cooperative courtship behaviour. However, field studies have revealed that dispersal can sometimes be high (Höglund & Alatalo 2014), with no clear indication that dispersal is counter-selected. In this context, a higher level of population viscosity could be maintained: (1) if dispersers preferentially join leks composed of close relatives, and/or (2) if dispersers incur sufficient reproductive costs after settling into a new lek consisting of non-relatives.

An elegant way to evaluate the reproductive costs of dispersal in the wild is by comparing ‘non-effective’ and ‘effective’ dispersal. Non-effective dispersal describes dispersal events that may or may not be followed by successful reproduction (Broquet & Petit 2009, Cayuela et al. 2018). This kind of dispersal is usually quantified by means of direct observations in the field: for instance, using capture–recapture methods (Lebreton et al. 2009, Lowe & Allendorf 2010). In contrast, effective dispersal describes dispersal events followed by successful reproduction and resulting in gene flow. Effective dispersal can be quantified using molecular approaches (Prugnolle & De Meeûs 2002, Broquet & Petit 2009). When no dispersal cost occurs or this cost is compensated (Cotto et al. 2014), one would expect similar non-effective and effective rates of dispersal, over a similar range of distance. In contrast, when dispersal entails immediate or delayed costs resulting in a loss of reproductive success, one would expect non-effective dispersal to exceed effective dispersal (Prugnolle & De Meeûs 2002, Broquet & Petit 2009).

In this study, we aimed to detect the footprint of kin selection and competition by examining the spatial structure of relatedness and by comparing non-effective and effective dispersal in a lekking bird with a cooperative breeding system, the Western capercaillie (*Tetrao urogallus*). We used capture-recapture and genetic data collected over a 6-year period (corresponding to two generations of capercaillie in our study system) in a spatially structured population of *T. urogallus* located in the Vosges Mountains of eastern France. Previous studies have highlighted a relatedness structure in males of this species at a small spatial scale, possibility driven by kin selection (Regnaut et al. 2006, Segelbacher et al. 2007). We hypothesized that male population viscosity is potentially maintained by two non-exclusive mechanisms: philopatry or high post-settlement costs, both of which are likely to enhance the genetic structure of a population. The first mechanism implies context-dependent dispersal, with males preferentially attending leks composed of relatives. At their first attendance at a lek, young males may preferentially join one consisting of relatives (likely their father’s) and then exhibit philopatric behaviour due to kin-selected benefits; alternatively, they may leave the lek (or attend another) to avoid kin competition, which should contribute to a decrease in population viscosity. The second mechanism allowing male population viscosity to be maintained could be high post-settlement costs due to intense intrasexual competition and the loss of kin-selected benefits. Given these hypotheses, we expected that non-effective dispersal would exceed effective dispersal (i.e. gene flow). We also examined female non-effective and effective dispersal, relatedness spatial structure and genetic structuring. We expected a lower relatedness structure in females than in males, likely due to inbreeding avoidance (Pusey 1987, Szulkin & Sheldon 2008) and the absence of kin-selected benefits (Höglund & Alatalo 2014). Young females may thus avoid leks composed of relatives, and older females may emigrate from leks composed of relatives. As female–female competition for mate acquisition is marginal in lek systems (Höglund & Alatalo 2014), we also expected limited post-settlement costs for females arriving to mate in a new lek; accordingly, we hypothesized that non-effective and effective dispersal for females would be relatively similar.

## Materials and methods

### Study area, sampling design and identification of individuals

The study was conducted on a spatially structured population of *T. urogallus* located in the Vosges Mountains of eastern France. Between 2010 and 2015, the collection of non-invasive samples (95 % faeces and 5 % feathers) allowed us to identify 109 individuals (61 males and 48 females) distributed among eleven leks (1 –18 individuals identified par year and per lek). A detailed description of the study area, the sampling method and the genotyping approach is provided in Appendix A. The Euclidean distance between leks ranged from 2.6 to 42 km, with a median of 18 km. The population was surveyed using the capture–recapture (CR) method over a 6-year period (2010-2015).

**Fig. 1.**
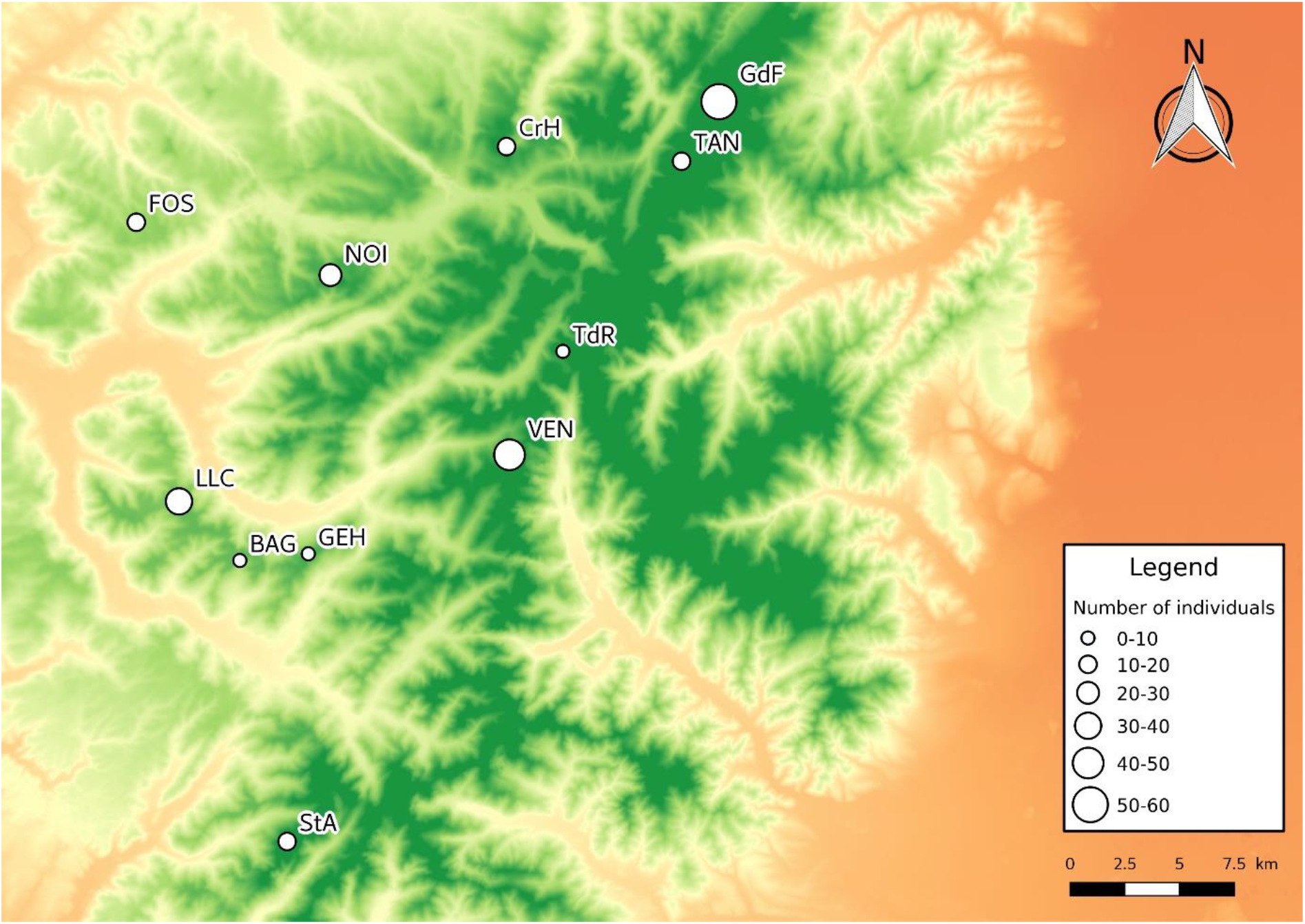
Map of the study area in the Vosges Mountains of eastern France. The 11 leks surveyed using capture–recapture and genetic methods are indicated according to size in terms of number of individuals. Each lek was identified by a three-letter code. The total number of individuals captured at least once over the 5-year study period was: GdF = 57, NOI = 26, CrH = 16, StA = 15, LLC = 31, GEH = 10, TdR = 5, BAG = 6, TAN = 13, FOS = 11, VEN = 46.

### Estimating non-effective dispersal from capture-recapture data

We quantified dispersal rates and distances for both sexes. Non-effective dispersal rate per generation and dispersal distance were estimated separately to facilitate model implementation. The models used in these analyses are described below. Prior to build capture-recapture models, we carried out Goodness-of-Fit tests (GOF). As no GOF test exists for multievent capture-recapture models, we used GOF tests designed for Cormack-Jolly-Seber models. We ran separate analyses for the two sexes. We examined the presence of transience and trap-dependence using respectively TEST3.SR and TEST2.CT implemented in the program UCARE (Choquet et al. 2009a). These analyses revealed the absence of transience in both females (overall test, df = 4, *χ*^2^ = 2.44, p = 0.65) and males (overall test, df = 4, *χ*^2^ = 1.22, p = 0.87). As well, we did not detect trap-dependence in females (overall test, df = 3, *χ*^2^ = 0.54, p = 0.91) and males (overall test, df = 3, *χ*^2^ = 0.52, p = 0.92). Therefore, we did not consider transience and trap-dependence in our capture-recapture models.

#### Quantifying the non-effective dispersal rate per generation

We modelled dispersal using multievent CR models (Pradel 2005); more specifically a model proposed by Lagrange et al. (2014) that allows the estimation of survival (ϕ) and dispersal (ψ) between numerous sites. By omitting site identity and distinguishing ‘individuals that stay’ from ‘individuals that move’, this model circumvents the computational issues usually encountered in standard multisite CR models when the number of sites is large. Lagrange’s model is based on states that embed information on whether an individual at *t* occupies the same site as it occupied at *t*−1 (‘S’ for ‘stay’) or not (‘M’ for ‘move’), as well as information about whether the individual was captured (‘+’) or not (‘o’) at *t*−1 and at *t*. We adapted this parameterization to estimate the proportion of dispersers (i.e. the individuals that changed lek at least once during their lifetime) per generation. The duration of the CR survey corresponded to approximately two western capercaillie generations in our population; the mean life expectancy derived from our survival estimates (see ‘Results’) was 3.16 years [ϕ/(1−ϕ), where ϕ is the survival probability]. Therefore, we can reasonably assume that the proportion of dispersers estimated in our model corresponds to the dispersal rate per generation. Note that we did not separate natal and breeding dispersal in our model as individual age cannot be assessed in our study system. Hence, fully resident individuals were assumed to be strictly philopatric; in contrast, dispersers may have moved before and/or after sexual maturity. This required extending Lagrange’s model to consider additional states that include information about fully resident behaviour (‘R’ for ‘fully resident’). In our model, an individual could belong to two alternative dispersal strategies during its lifetime: either a disperser that may be in state ‘S’ or ‘M’ depending on its movement status between *t−1* and *t*; or a fully resident individual that always remained in the same lek (‘R’). This led to the consideration of 10 states and 4 events (Appendix B, Table B1), which were coded in an individual’s capture history and reflect the information available to the observer at the time of capture.

To quantify the proportion of dispersers per generation, we coded the initial states of the model in two steps (Fig. 2): the first step embedded the probability μ that an individual is assigned to the disperser strategy; 1−μ corresponds to a probability of belonging the fully resident strategy. The second step included the movement and capture state of a newly encountered individual (Fig. 2). When captured for the first time, a disperser could be in state oS+ or oM+. Yet as these two states cannot be distinguished in Lagrange’s model or its extensions (Lagrange et al. 2014; Cayuela et al. 2017, 2018), we arbitrarily kept the state oM+ (Fig. 2). Fully resident individuals were assigned the state oR+. We then considered three modelling steps in which the information of the state descriptor was progressively updated in the model: survival (ϕ), dispersal (ψ) and recapture (*p*). Each step was conditional on all previous steps. As is standard in a multievent model framework (Pradel 2005, Lagrange et al. 2014), in the transition matrix, the rows correspond to time *t*−1, the columns to time *t*, and whenever a status element is updated to its situation at *t*, it is shown in bold and stays bold throughout the following steps. In the first step, survival information was updated: an individual could survive with a probability of ϕ or could die (**d**) with a probability of 1−ϕ. This resulted in a transition matrix with 10 states of departure and 7 intermediate states of arrival (Fig. 2). In the second modelling step, dispersal was updated. An individual belonging to the disperser strategy could move (**M**) from the lek it occupied with a probability of ψ or could stay (**S**) with a probability of 1−ψ. An individual belonging the fully resident strategy always stayed in the same lek; its probability of staying in the same lek was fixed at 1. This led to a matrix of 7 states of departure and 7 states of arrival (Fig. 2). In the third step, the recapture information was updated: an individual could be recaptured with a probability of *p* or not with a probability of 1−*p*.

**Fig. 2.**
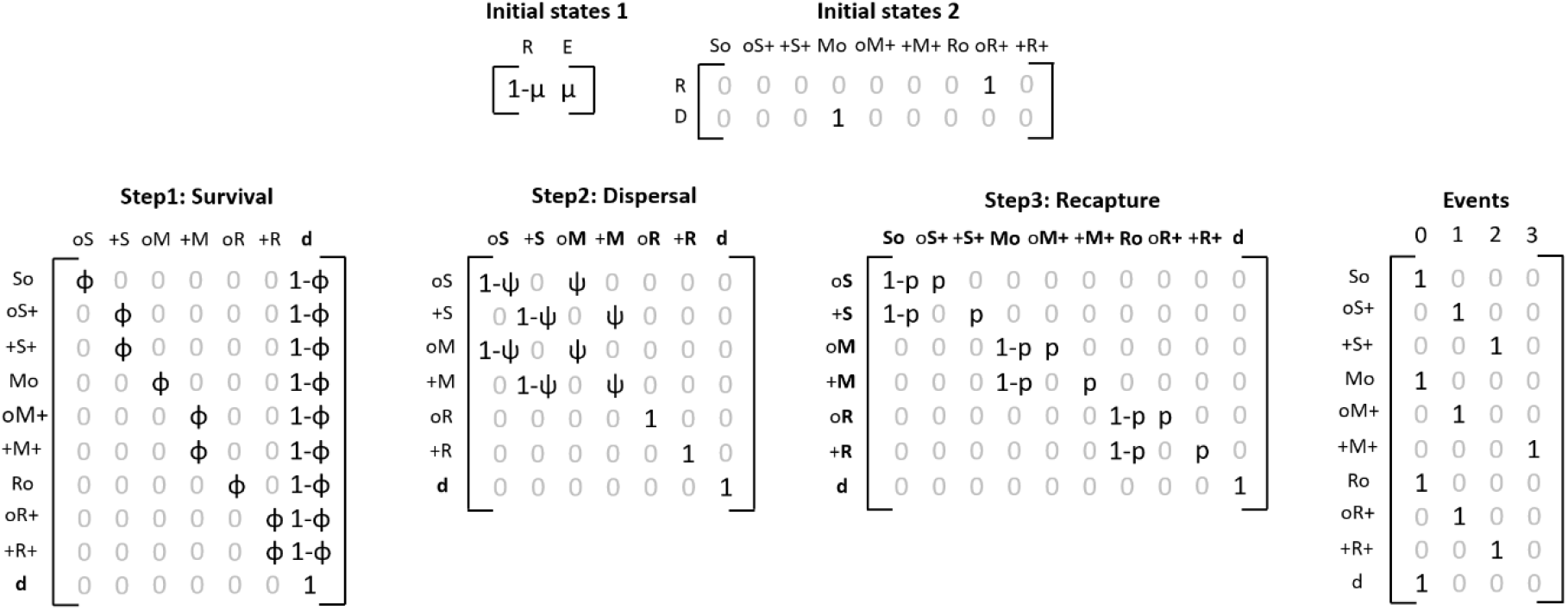
The CR multievent model with its state-transition and event matrices for quantifying the proportion of non-effective dispersers per generation. The initial states were coded in two steps (Initial states 1 and 2). In initial state 1: μ = proportion of dispersers per generation, E = dispersers, R = fully resident individuals. The state–state transitions were then modelled through three steps: survival ϕ, dispersal ψ, and recapture *p*. In the transition matrix, the rows correspond to time *t*−1, the columns to time *t*; whenever a status element is updated to its situation at *t*, it is shown in bold and stays bold throughout the following steps. For a detailed description of states and events, see Appendix B, Table B1.

This parameterization was implemented in the E-surge program (Choquet et al. 2009b). Competing models were ranked through a model-selection procedure using Akaike information criteria adjusted for a small sample size (AICc) and AICc weights. If the AICc weight of the best-supported model was less than 0.90, we performed a model-averaging procedure. Our hypotheses concerning survival, dispersal and recapture were examined using the general model [μ(SEX), ϕ(SEX), ψ(SEX), *p*(SEX + YEAR)], in which two effects were considered: a sex effect (SEX), included in the model as a discrete covariate; and a year effect (YEAR). From this general model, we tested all possible combinations of effects and ran 32 competing models (Appendix C, Table C1).

#### Quantifying annual non-effective dispersal rates and distances

In a recent paper, Tournier et al. (2017) extended Lagrange’s model by breaking down dispersal (ψ) into the distinct parameters of departure (τ) and arrival (α). This new parameterization allows the quantification of the proportion of individuals arriving in sites with different characteristics or located at different distances from the source site. We adapted this parameterization for the needs of our study to consider states incorporating information about the capture of an individual (‘+’ or ‘o’) at *t*−1 and *t* and its movement status. We also included information about the Euclidian distance covered by dispersers between a departure and an arrival patch. This was incorporated in the model using five Euclidean distance classes (noted ‘1’ to ‘5’) – to date, Euclidean distance cannot be included as a continuous variable in multistate and multievent capture-recapture models. The distance classes were fixed according to the movement distances recorded in the population, ranging from less than 1 km to 29 km: 0–6 km (‘1’), 6–12 km (‘2’), 12–18 km (‘3’), 18–24 km (‘4’) and 24–30 km (‘5’). The number of distances classes was limited to five to facilitate model implementation; the number of model states increases with the number of classes. This led to the consideration of 19 states and 8 events (Appendix B, Table B2).

When captured for the first time, the state of an individual could be oS+ or oM+. Yet as these two states cannot be distinguished in Lagrange’s model or its extensions (Lagrange et al. 2014, Cayuela et al. 2017), we arbitrarily kept the state oS+ (Fig. 3). We then considered four modelling steps: survival (ϕ), departure (τ), arrival (α), and recapture (*p*). In the first step, the information about survival was updated. An individual could survive with a probability of ϕ or could die (**d**) with a probability of 1−ϕ. This led to a matrix with 19 states of departure and 5 intermediate states of arrival (Fig. 3). In the second step, departure was updated. An individual could move (**M**) from the lek it occupied with a probability of τ or could stay (**S**) with a probability of 1−τ. This led to a matrix of 5 states of departure and 5 states of arrival (Fig. 3). In the third step, we updated the information about arrival. An individual that moved could arrive in a patch located in the first four distance classes (**1**, **2**, **3** or **4**) from the source patch with a probability of α1, α2, α3 or α4, or could arrive in a lek located in distance class **5** with a probability of 1−(α1+α2+α3+α4). This led to a matrix with 5 states of departure and 13 states of arrival (Fig. 3). In the fourth step, recapture was updated (Fig. 3). An individual could be recaptured with a probability of *p* or not with a probability of 1−*p*, resulting in a transition matrix with 13 states of departure and 19 states of arrival. The last component of the model linked events to states. In this specific situation, each state corresponded to only one possible event (Fig. 3). The information about individuals’ dispersal status, distance, and capture is coded in the events (Appendix B, Table B2).

**Fig. 3.**
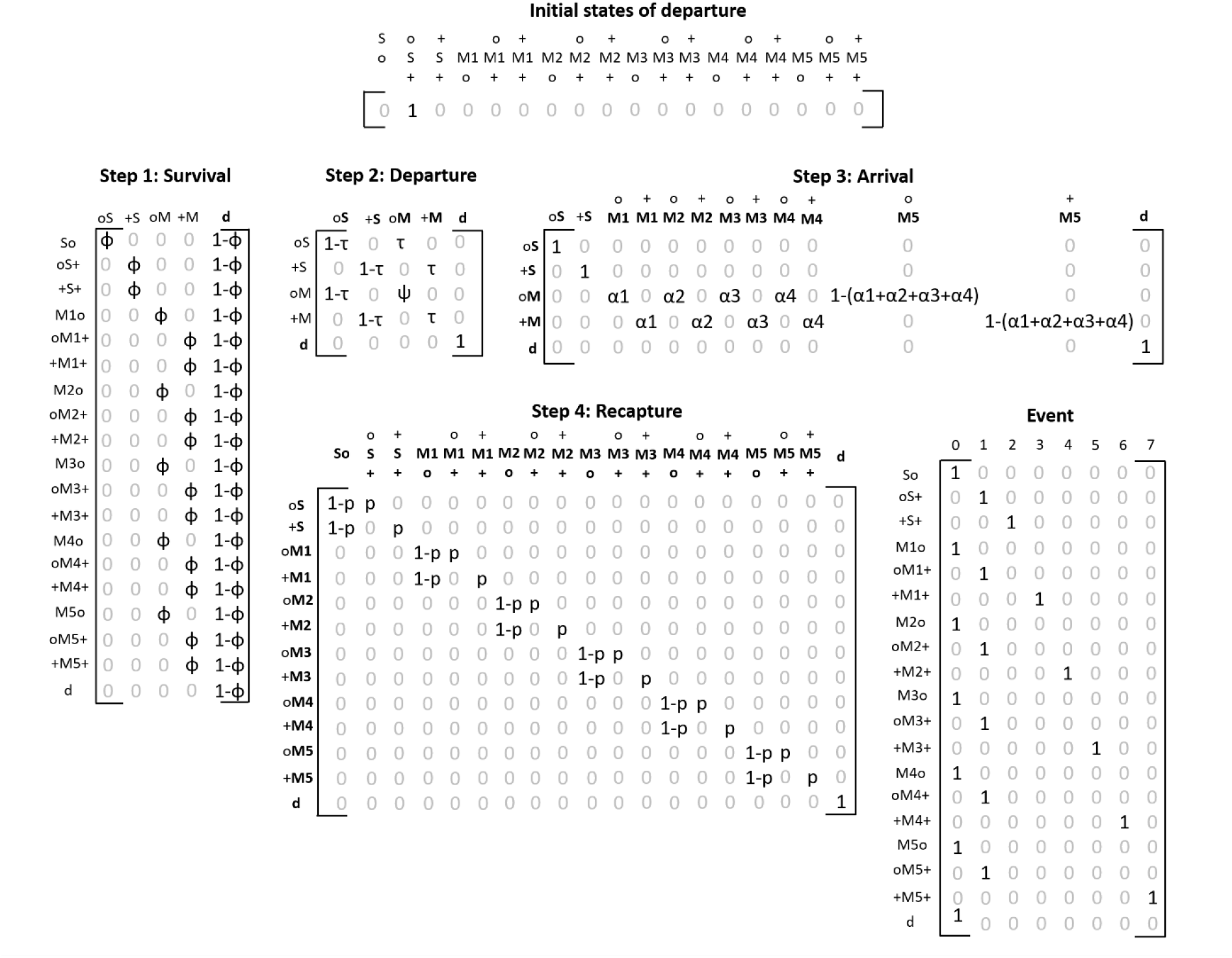
The CR multievent model with its state-transition and event matrices for quantifying annual non-effective dispersal rates and distances. The initial states were coded in a single step. The state–state transitions were then modelled through four steps: survival ϕ, departure τ, arrival a and recapture *p*. In the transition matrix, the rows correspond to time t−1, the columns to time *t*, and whenever a status element is updated to its situation at *t*, it *i*.

As for the previous model, this parameterization was implemented in the E-surge program. Competing models were ranked through a model-selection procedure (provided in Appendix C) using Akaike information criteria adjusted for a small sample size (AICc) and AICc weights. We performed model averaging if the AICc weight of the best-supported model was less than 0.90. We built a model-selection procedure for each type of distance (i.e. Euclidean, topographic and ridge). Our hypotheses concerning state-state transition probabilities were tested using the general model [ϕ(SEX), τ(SEX), α(5 + SEX), *p*(SEX + YEAR)]. From the general model, we tested all the possible combinations of effects and built 32 competitive models (Appendix C, Table C1).

### Assessing spatial genetic structure and relatedness

We examined the spatial genetic structure and relatedness for both sexes. A total of 92 individuals were kept for the population genetics analyses. We used 11 microsatellite markers for our analyses (ADL184, ADL230, BG15, BG16, BG18, LEI098, TuT1, TuT2, TuT3, TuT4, ADL142; see Appendix A). We used genetic approaches designed to detect contemporary genetic structure and relatedness pattern. The Bayesian clustering approach (STRUCTURE) used in our analyses is designed to detect F0 immigrants, and their offspring F1 and eventually F2 (Pritchard et al. 2000, Corander et al. 2003, Broquet & Petit 2009). Parentage analyses typically describe the genealogical relationships (i.e. relatedness level) between individuals through successive generation and applies to recent times (Wang 2004, 2012; Städele & Vigilant 2016). Furthermore, the use of individual-based approaches show less inertia than approaches (e.g. *F_st_*-based methods) assessing the evolution of allele frequencies through many generations and are therefore more suitable to analyse contemporary gene flow (Landguth et al. 2010). Hence, we can assume that non-effective dispersal, relatedness structure and gene flow are inferred at relatively similar (and recent) time scale.

#### Examining genetic structure

Evidence for scoring errors, large allele dropout, presence of null alleles, departure from Hardy–Weinberg equilibrium (HWE) and linkage disequilibrium (LD) between all pairs of loci were assessed with Micro-Checker v2.2.3 and ARLEQUIN v3.5.1.3 (Van Oosterhout et al. 2004, Girard & Angers 2008, Excoffier et al. 2005). These tests were conducted only on the four large leks containing at least ten individuals (NOI = 10, LLC = 12, VEN = 20 and GdF = 26) with the Bonferroni correction for multiple comparisons (P<0.05; Rice 1989). Analyses of the spatial genetic structure were performed for all individuals, for males only and for females only with Bayesian clustering of genotypic data, implemented in STRUCTURE v2.3.3 (Pritchard et al. 2000), to estimate the most likely number of homogeneous genetic clusters (Appendix D). The Markov chain Monte Carlo (MCMC) algorithm was run using the admixture model with correlated allele frequencies (Falush et al. 2003) for 500,000 steps after 2,000,000 initial burn-in steps, with no a priori information on an individual’s lek. Ten independent runs were performed for each value of K, from 1 to 10. The most likely number of genetic clusters in the dataset was suggested with the deltaK method (Evanno et al. 2005), which was applied using tools available from the STRUCTURE HARVESTER website (Earl & von Holdt 2012). However, the deltaK method is not appropriate when the true K = 1, so we first tested for this possibility by examining whether lnP(D) (an estimate of the posterior probability of the data for a given K) was maximum for K = 1. We also verified that the different runs for each K were similar using the similarity coefficients (H’ values) of the software CLUMPP v 1.1.1 (Jakobsson and Rosenberg 2007). The Greedy algorithm was used for K = 1 to 5, then the LargeKGreedy algorithm was used for K = 6 to 10. To confirm that sex-specific genetic structure (see ‘Results’) was linked to higher genetic proximity of males within leks, we first compared male and female genetic structure with a discriminant analysis of principal components (DAPC) using leks as a priori (DAPC; Jombart et al. 2010). The DAPCs were computed with the adegenet package in R (Jombart 2008).

#### Examining relatedness structure

We examined relatedness structure by comparing intra- and inter-lek relatedness (*r*xy) for males and females using COANCESTRY v1.0.1.8 (Wang 2011). Simulations were performed to identify the best relatedness estimator; these consisted of 600 dyads spread equally across six categories of relatedness: parent–offspring (*r*xy = 0.5), full siblings (*r*xy = 0.5), half siblings/avuncular/grandparent–grandchild (*r*xy = 0.25), first cousins (*r*xy = 0.125), second cousins (*r*xy = 0.03125), and unrelated (*r*xy = 0). In addition, we used a permutation approach to test if the average relatedness of an individual was significantly higher within the lek of its first capture than a randomly chosen lek. This permutation approach was performed separately for males and females in the goal to compare pattern between sex. We first computed the average relatedness of all individuals to each lek (based on the males present in the lek and excluding the individual if present). For each individual, we then randomly chose a lek and created a vector of average relatedness of the permutated ‘first capture’ lek. The mean of this vector was calculated, and a distribution of this statistic was drawn based on 1,000 iterations and compared with what we observed in our original dataset. The *P*-value was calculated by dividing the number of values in the distribution that were higher or equal to the original dataset with the number of permutations (i.e. 1,000). The same approach was then used with departure from a lek to test if average relatedness might have an impact on the decision of an individual to emigrate from a given lek and compare pattern between sex.

### Estimating effective dispersal from genetic data

We assessed effective dispersal (i.e. gene flow) distance in males and females, notably in response to various landscape features (topographic distance, slope and ridge) likely to affect dispersal.

#### Inter-individual measures of genetic differentiation

The use of autosomal nuclear markers does not prevent the detection of sex-specific differences in dispersal as long as only adults are sampled (Goudet et al. 2002). All the genetic analyses were thus performed separately on males (dataset M) and females (dataset F). For each dataset, we computed two kinds of inter-individual pairwise genetic distances: the Bray-Curtis percentage dissimilarity metric (Bc; Legendre & Legendre 1998) and the dissimilarity metric based on the proportion of shared alleles at each locus (Dps). In each dataset, both metrics were highly correlated (Pearson’s correlation coefficient > 0.90).

#### Statistical analyses

For each dataset (M and F) and each measure of genetic differentiation GD (Bc and Dps), we first performed a spatial autocorrelation analysis with a non-directional Mantel correlogram (Smouse & Peakall 1999) using the MATLAB software-coding environment (Mathworks, Inc.) to determine the spatial scale of indirect gene flow (Anderson et al. 2010). For this purpose, Euclidean distance classes were defined every 5000 m (up to 40 km), resulting in 8 binary matrices representing the membership of individual pairs to the distance class tested (with ‘0’ for pairs of individuals belonging to the same distance class and ‘1’ otherwise). Each binary matrix was compared to the Bc or the Dps matrix using a simple Mantel test with 1,000 permutations. We then plotted Mantel correlation values over distance classes, with a 95% confidence interval determined by the random removal of 20% of individuals (1,000 iterations; Peterman et al. 2014).

As dispersal may be affected by functional landscape connectivity, we also assessed the spatial scale at which various landscape features likely to affect movement in *T. urogallus* might explain variance in sex-specific measures of genetic differentiation (Keller et al. 2013). In addition to Euclidean distance (ED), we considered three kinds of landscape predictors (see Appendix E for details): topographic distance (*Dtopo*) (assuming that individuals may not fly in a straight line, but may follow the topographic relief), effective distance along slopes (*Dslope*) (assuming that individuals may avoid steep zones) and effective distance along ridges (*Dridge*) (assuming that individuals may prefer ridge zones). For each measure of genetic differentiation GD (Bc and Dps) and for each sex, we designed a complete model as follows: *GD* ~*ED* + *Dtopo* + *Dslope* + *Dridge*. We did not run this model on all possible pairs of individuals, but selected various subsets of pairwise data by defining a maximum Euclidean distance of S between sample points. S ranged from 14,000 m (a distance chosen so that no individual was excluded from the neighbouring graph; Jombart et al. 2008) to 46,000 m (the maximum Euclidean distance between two individuals) in 1000 m increments. For each subset, we ran the full model in a standard multiple linear regression and plotted the model fit R^2^ against S to identify, for each sex, the spatial scale that optimized the amount of variance explained in each metric of genetic differentiation.

## Results

### Estimating non-effective dispersal

Concerning the proportion of dispersers per generation, the best-supported CR model was [μ(.), ϕ(.), ψ(.), *p*(sex)] (Appendix C, Table C1). However, as the AICc difference with the second-ranked model was small (ΔAICc = 0.41), we performed model averaging (from the complete set of models). Model-averaged estimates of recapture probabilities indicated that males (*p* = 0.72±0.06) were more detectable than females (*p* = 0.46±0.06). Survival did not differ substantially between the two sexes; the survival probability of males (ϕ = 0.75±0.04) was only marginally lower than that of females (ϕ = 0.79±0.06). The proportion of dispersers per generation (μ) was also relatively similar in both sexes; 0.38±0.09 in females and 0.30±0.15 in males. In the individuals with a disperser strategy, the annual dispersal probability ψ did not differ between sexes; it was 0.67±0.15 in females and 0.64±0.15 in males.

Concerning annual non-effective dispersal rates and distances, the best-supported CR model was [ϕ(.), τ(sex), α(5), *p*(sex)] (Appendix C, Table C2). Again, as the AICc difference with the second-ranked model was small (ΔAICc = 0.05), we performed model averaging (from the complete set of models). The annual departure rate slightly differed between sexes: females had very marginally higher dispersal probability (τ = 0.23±0.15) than males (τ = 0.16±0.07). Note that departure rate showed little variation between years (see Appendix C). Arrival probability was similar between sexes and decreased with the Euclidean distance between leks (Fig. 6). The probability of arriving in a lek located at a distance ranging from 0 to 6 km was 0.49±0.12 in females and 0.50±0.12 in males. At distances between 6 and 24 km, the arrival probability was relatively constant (ranging from 0.11 to 0.17) in both sexes. At distances from 24 to 30 km, arrival probability substantially decreased (0.03±0.03 in females and 0.07±0.07 in males).

### Assessing spatial genetic structure and relatedness

We found no evidence of genotyping errors, large allele dropout or null alleles. No significant deviation from the Hardy-Weinberg equilibrium (HWE) was observed in three of the four large leks; linkage disequilibrium (LD) was observed between the loci ADL230 and BG15 for the lek NOI after Bonferonni correction (P < 0.05). Because the small sample size could be linked to this LD, we kept these loci for further analyses and thus consider our molecular marker dataset as neutral and biparentally inherited.

When considering all individuals, the optimal number of genetic clusters found with the STRUCTURE software was two (Appendix D, Fig. D1). However, the optimal number of genetic clusters for males alone was three, and for females it was one (Fig. 4; Appendix D, Fig. D2 and D3). The DAPCs suggest higher male genetic proximity inside a given lek in comparison to females (Appendix D, Fig. D4 and D5).

Concerning relatedness analyses, according to our simulations, the best *r*xy estimator was LynchRD, which has a similar mean *r*xy in comparison to true values (0.22 versus 0.23) and a correlation of 0.61 (Appendix D, Table D1). Based on this estimator, males and females differed in terms of relatedness. Males had a significantly higher intra-lek average relatedness than females (t=5.10, df=254.55, *P* < 0.001; Fig. 5). Both males and females had significantly higher average relatedness within the lek of their first capture (*P* < 0.001 for both), but only females showed significantly higher average relatedness in leks from which they emigrated (*P* = 0.042; Appendix F, Fig. F1).

**Fig. 4.**
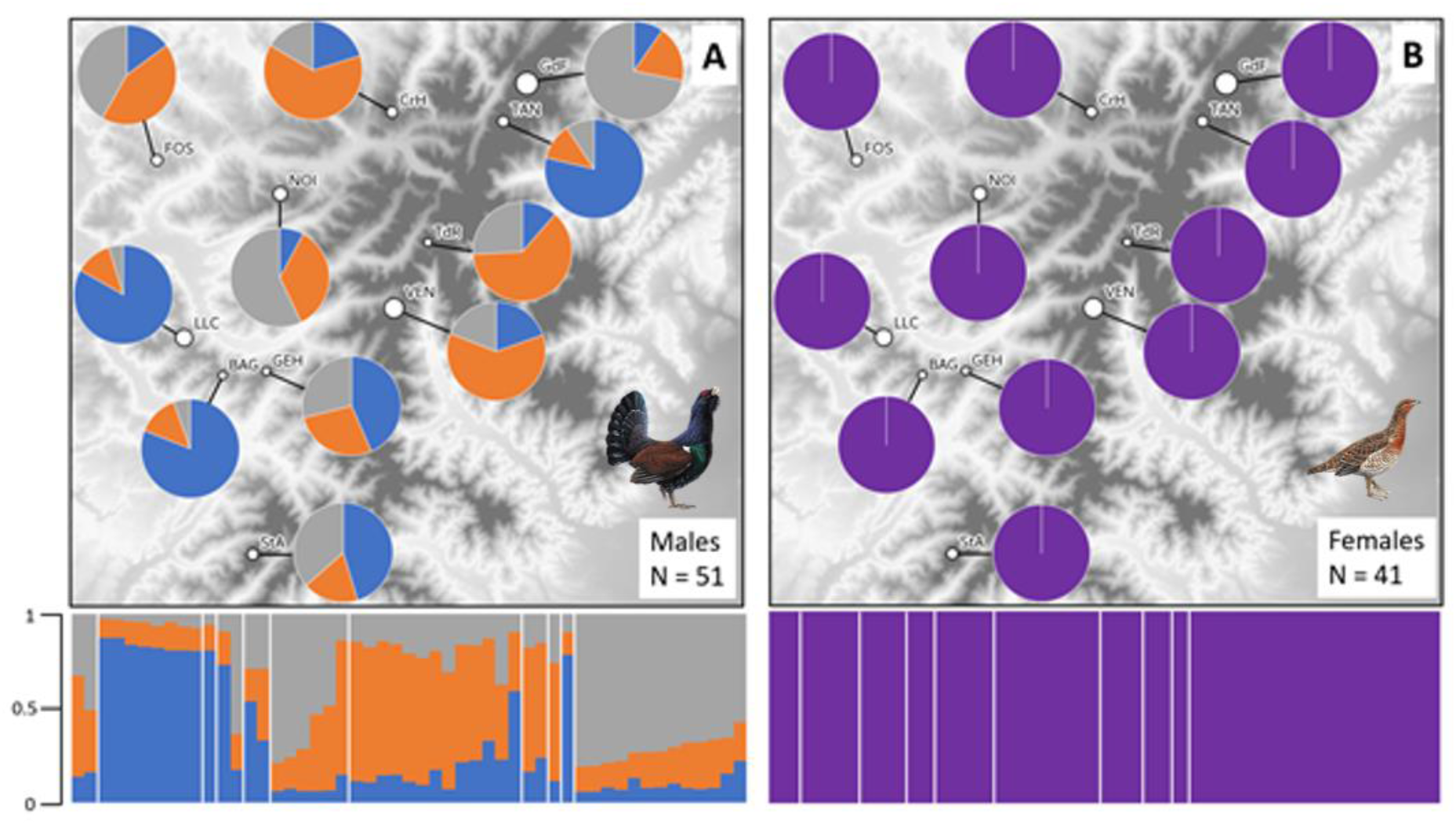
Outputs from STRUCTURE software showing high population genetic structure in males (A: K = 3) and the absence of genetic structure in females (B: K = 1). The sample size was *n* = 51 in males and *n* = 41 in females.

**Fig. 5.**
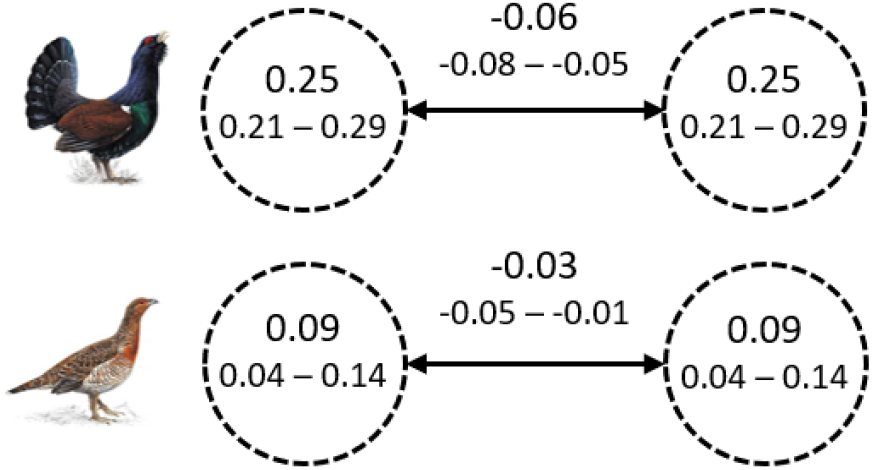
Spatial structure of relatedness in the study area population of T. urogallus (top: males; bottom: females) The mean relatedness coefficients r were calculated within leks (circles) and between leks (arrows) and are provided with their 95% CI. The sample size was *n* = 51 in males and *n* = 41 in females.

**Fig. 6.**
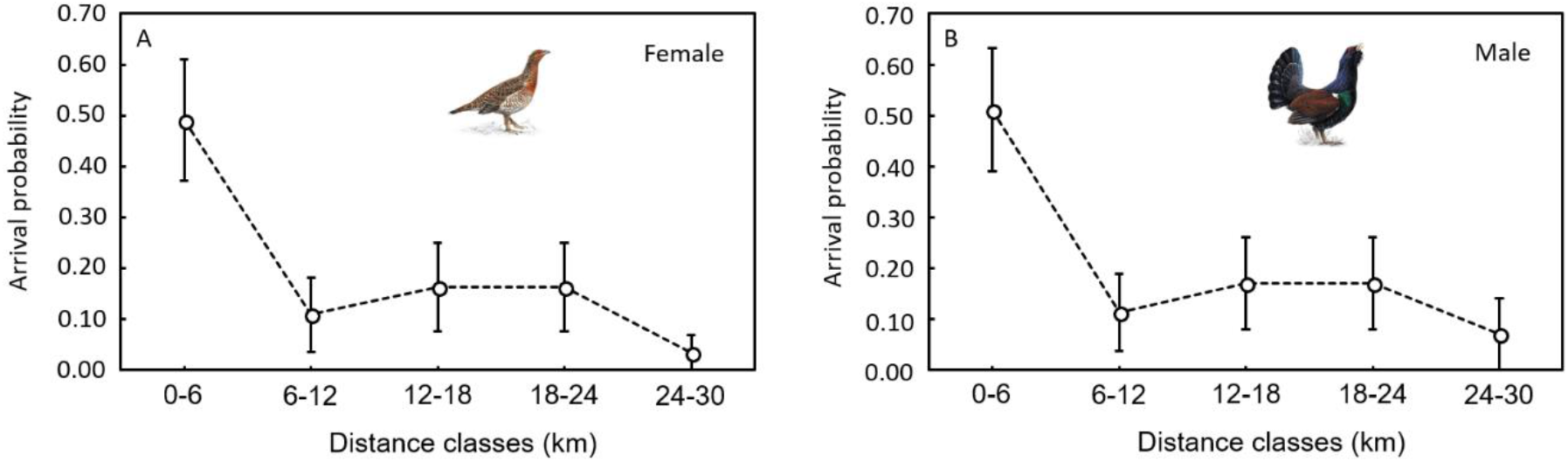
Annual non-effective dispersal distance in T. urogallus [left: females (n = 48), right: males (n= 61)]. The arrival probabilities (with standard errors shown in error bars) were model averaged.

### Estimating effective dispersal distance

Autocorrelograms indicated spatial patterns of isolation by distance in both sexes, with significant positive spatial autocorrelation occurring up to 10 km in both males and females (Fig. 7 A and B). Nevertheless, females showed positive (though non-significant) spatial autocorrelation up to 20 km, whereas males showed significant negative autocorrelation from 15 km, suggesting higher effective dispersal distances in females than in males. Patterns of isolation by distance were very similar when considering Dps genetic distances.

**Fig. 7.**
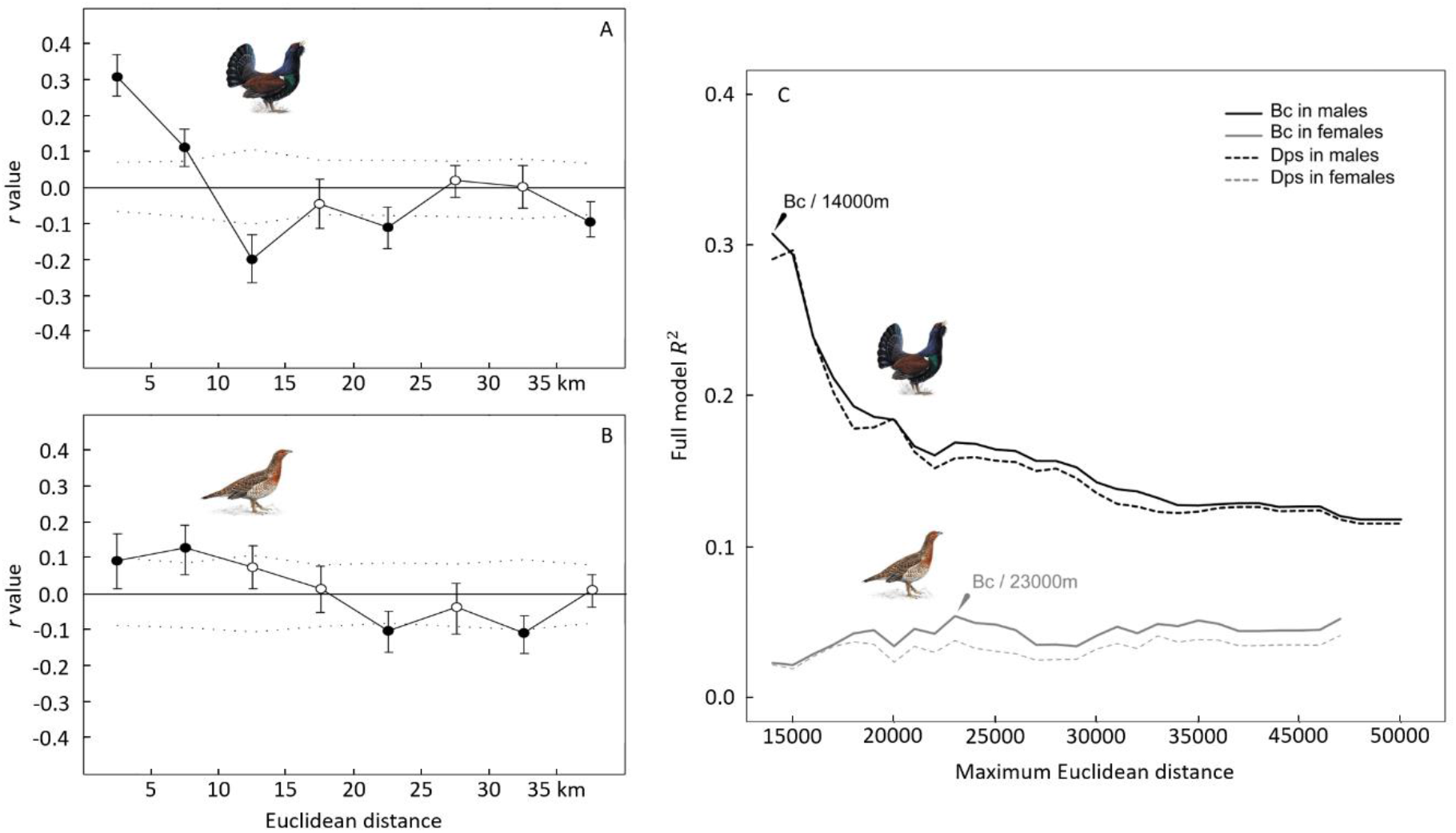
Effective dispersal distances in T. urogallus. A and B: Mantel correlograms showing the relationships between Bc genetic distances and Euclidean distance classes (defined every 5000 m) in males (A) and females (B). The dots indicate the value of r: standard Mantel correlation with 1,000 permutations (black circles, pvalue < 0.05). Error bars bound the 95% CI of r as determined by the random removal of 20% of populations. Upper and lower confidence limits (dotted lines) bound the 95% CI of the null hypothesis of no spatial structure as determined by permutations. C: Plot of model fit against maximum pairwise Euclidean distance in males (black) and females (grey). Solid lines: model fit with Bc dissimilarity metric as the dependent variable. Dotted lines: model fit with Dps dissimilarity metric as the dependent variable. Arrows indicate the best combination of maximum Euclidean distance and genetic distance for each sex. The sample size was 51 in males and *n* = 41 in females.

The plot of model fit R^2^ against maximum pairwise Euclidean distances indicated that model fit was always higher in males (R^2^ ranging from 0.115 to 0.308) than in females (R^2^ ranging from 0.019 to 0.054), whatever the Euclidean distance and the considered measure of genetic differentiation (Fig. 7C). This did not result from unbalanced sample size in males (*n* = 51) and females (*n* = 41). Furthermore, model fit values were always higher (or at least equal) when using Bc rather than Dps, in both sexes. In males, the optimal maximum Euclidean distance was 14,000 m (R^2^ = 0.308), whereas it was 23,000 m in females (R^2^ = 0.054). These maximum Euclidean distances were in line with the autocorrelograms and roughly corresponded to the first distance class associated with negative autocorrelation in each sex. Note that, in both sexes, the main contributor to the variance in measures of genetic differentiation was the topographic distance, indicating that topographic relief is a better predictor of effective dispersal than Euclidean distance in this species (see Appendix E for details).

### Comparing non-effective and effective dispersal patterns

Overall, our results showed that non-effective and effective dispersal patterns differ between sexes. In females, the effective dispersal distances were highly congruent with the non-effective dispersal distances. The proportion of female dispersers settling in a lek located at a distance of more than 24 km was dramatically lower (<10%) (Fig. 4), which is in accordance with the upper estimate of gene flow distance (23 km) (Fig. 7). In males, the non-effective dispersal distances exceeded effective dispersal distances. The pattern of male non-effective dispersal distance was relatively similar to that found in females, with substantially fewer dispersers beyond a distance of 24 km (Fig. 4). Yet the maximum gene flow distance was 14 km (Fig. 7), resulting in a 10-km gap between non-effective and effective dispersal distances in males.

## Discussion

Our findings revealed stronger spatial structure of relatedness in males than in females, despite that males disperse at a similar rate and distance than females. They also indicated that the population viscosity allowed male cooperation through two non-exclusive mechanisms. First, the results showed that at their first lek attendance, males aggregated in a lek composed of relatives and tended not to leave this patch in order to avoid kin competition. Second, non-effective dispersal distance dramatically outweighed effective dispersal distance, which suggests that dispersers incurred high post-settlement costs. These two mechanisms result in strong population genetic structure in males. In females, our study revealed a lower spatial structure of relatedness compared to males. At their first lek attendance, females preferentially arrived in a lek composed of relatives. Yet they also preferentially tended to emigrate from this type of lek, likely due to inbreeding risk, which weakens the spatial structure of relatedness. Our results also found that non-effective dispersal and effective dispersal distances were highly similar in females, which suggests the absence of post-settlement costs. So both this and inbreeding avoidance are likely to explain the low genetic structure in females.

### Non-effective dispersal rate is high and does not differ between sexes

Our study revealed that non-effective dispersal rates per generation and annual dispersal probability were relatively similar for both sexes. Overall, non-effective dispersal between leks was high in our study area: 38% of females and 30% of males dispersed at least once during their lifetime. The annual non-effective dispersal rates were also high: 23% in females and 16% in males. This result goes against the conventional view that *T. urogallus* males are highly philopatric (Höglund & Alatalo 2014). Yet to our knowledge, our study is the first to use long-term CR data to quantify dispersal rates in this species. In addition, while previous studies have mainly focused on movements at a relatively small spatial scale (Hjeljord et al. 2000, Wegge et al. 2007), the large spatial extent of our study allowed the detection of long-distance dispersal events (max. = 29 km). Although our sample size is relatively small (*n* = 107), we are confident that the patterns drawn in our study are reliable. If a difference of dispersal rate (or distance) actually exists between sexes, it should be small.

### Females likely avoid inbreeding and do not pay high dispersal costs

Our results highlighted that the difference between intra-lek and inter-lek relatedness was relatively weak in females. Within leks, the mean coefficient of relatedness was 0.09, which corresponds to genealogical levels of first and second cousins; between leks, the mean coefficient was −0.06, indicating that females are only weakly related to the lekking males. The findings indicate that, at first attendance in a lek, females preferentially arrive in a lek composed of close relatives. Yet our study revealed that 38% of females change lek at least once during their lifetime; dispersing females have a 64% chance of dispersing each year. We found that females preferentially emigrate from leks of males that are close relatives (Appendix F, Fig. F1), likely to avoid inbreeding risks. A similar inbreeding-avoidance mechanism was highlighted by Lebigre et al. (2010) in the closely related species *Tetrao tetrix*. Furthermore, our results showed that non-effective dispersal distances for females (less than 5% of dispersers move farther than 24 km) are highly congruent with effective dispersal distances (the maximum gene flow distance is 23 km). This suggests the absence of substantial reproductive costs after settling into a new lek, likely due to reduced female–female competition for mate acquisition in lekking birds (Höglund & Alatalo 2014). In females, inbreeding avoidance behaviour and the absence of high post-settlement costs likely reduce the spatial structure of relatedness, as well as the population genetic structure.

### Context-dependent dispersal and high post-settlement costs facilitate cooperative courtship in males

In contrast, in males, the results revealed the existence of a strong spatial structure of relatedness. Within leks, the mean coefficient of relatedness was 0.25, which corresponds to genealogical levels equal to half sibling, avuncular or grandparent–grandchild. Between leks, the mean coefficient was −0.06, indicating that males are unrelated. This pattern is congruent with those found in two previous studies on *T. urogallus* (Regnaut et al. 2006, Segelbacher et al. 2007) and indicates that males preferentially join leks composed of relatives. Our study revealed that this spatial structure of relatedness is maintained despite relatively high non-effective dispersal: 30% of the males disperse at least once during their lifetime, and these males have a high propensity to disperse each year (0.65). These findings suggest that the maintenance of male population viscosity is based on two non-exclusive mechanisms. First, after brood dissolution, young males tend to aggregate in leks composed of close relatives (likely their father’s). When they emigrate from a lek, emigration does not seem to be associated with the social context within the lek of departure; permutation analyses indicate that relatedness does affect emigration. The high capture probability of males (0.72) does not suggest that we failed to detect an effect due to a methodological bias. Yet it should be kept in mind that the number of dispersal events was limited in our dataset, which likely limited our ability to detect a signal. Second, the findings suggest that male population viscosity might also be maintained by high dispersal costs. Non-effective dispersal distance dramatically outweighed effective dispersal distance, suggesting that dispersers may pay acute post-settlement costs. The fitness loss after settling into a new lek composed of non-relatives could be caused by intense male–male competition, associated with territorial and agonistic behaviour (Höglund & Alatalo 2014) and/or the loss of kin-selected benefits (Krakauer 2005, DuVal 2007). However, although these results are highly congruent with predictions of theoretical models (El Mouden & Gardner 2008), we are aware that the evidence is indirect; these conclusions would need to be confirmed by pedigree analyses (which were not feasible in our study due to the low number of neutral markers) and/or experimental studies.

In summary, our results strongly suggest that male aggregation in leks composed of relatives and enhanced dispersal costs could maintain population viscosity, creating the conditions conducive to cooperative courtship. While cooperative males are more likely to benefit kin in viscous populations via inclusive fitness, they must also compete for limited mating opportunities with these same kin (Platt & Bever 2009). As a result, kin competition could cancel out to some degree the benefits of kin cooperation, possibly promoting kin-dependent dispersal in lek systems. However, in our study, we failed to detect an effect of kin competition on departure probability. The strong spatial structure of relatedness indicates that, if dispersal driven by kin competition exists, its consequences on lek social composition is weak. This might be due in part to the fact that the recent decline of the population in our study area (possibly caused by anthropogenic habitat degradation) has resulted in a marked decrease in lek size, which may limit the risk of kin competition.

### Role of direct and indirect benefits in lek system evolution

Previous studies reported contradictory results about the role kin selection in the evolution of bird lek systems. Several studies in manakins (Loiselle et al. 2006, McDonald 2009, Davis et al. 2015), bowerbirds (Madden et al. 2004) and grouses (Gibson et al. 2005, Lebigre et al. 2008, Bush et al. 2010) did not detect any increase of relatedness level within leks and concluded that the maintenance of male aggregations could mostly be due to direct rather than indirect fitness benefits. Three alternative hypotheses have been proposed: the hot-spot hypothesis states that males are hypothesized to sequentially cluster in areas of high female density or movement (Bradbury & Gibson 1983); the hotshot hypothesis postulates that subordinate males settle near dominant males with high reproductive success (Beehler & Foster 1988); in the delayed benefits hypothesis, subordinate males receive direct-fitness benefits later in life when they replace higher ranking males on the leks (McDonald & Potts 1994, Kokko & Johnston 1999). To date, the role of these different mechanisms in lek system evolution is still debated (Loiselle et al. 2006, Höglund & Alatalo 2014). Other studies, as ours, detected an increase of the relatedness level within leks in manakins (Shorey et al. 2000, Höglund & Shorey 2003, Francisco et al. 2009, Concannon et al. 2012), bowerbirds (Reynolds et al. 2009), peafowls (Petrie et al. 1999), wild turkeys (Krakauer 2005), and grouses (Höglund et al. 1999, Regnaut et al. 2006). These studied defended the hypothesis that kin selection and indirect fitness benefits play a central role in the evolution of cooperative breeding systems. They stated that population viscosity can be maintained by male philopatry and kin-recognition mechanism including self referent phenotypic matching (Petrie et al. 1999). In this context, our study also suggests that males dispersing to leks composed of non-relatives may incur high post-settlement costs due to the loss of kin-selected benefits. To date, the role of direct and indirect benefits in male aggregation and lek system formation still remains controversial. Broader analyses should be undertaken to quantify within-lek relatedness level in a greater number of bird species, and population per species, to draw general conclusions about the mechanisms shaping lek system evolution.

## Acknowledgements

The genetic monitoring of capercaillie in the Vosges Mountains was funded by the Life+ Project “Des Forêts pour le Grand tetras”, by the Natura2000 network and by the regional programme of the Capercaillie National Action Plan initiated by the French Ministry of Environment. The project largely relied on the work of volunteers who collected samples during the six years of the study: Antoine Andre, Didier Arseguel, Samuel Audinot, Alix Badre, Etienne Barbier, Dominique Becker, Bernard Binetruy, Frédéric Bocquenet, Noémie Castaing, Sebastien Coulette, Stéphane Damervalle, Luc Dauphin, Richard Delaunay, Lucile Demaret, Michel Despoulin, Laurent Domergue, Vincent Drillon, Christian Dronneau, Fabien Dupont, Arnaud Foltzer, Patrick Foltzer, Marc Gehin, Cyril Gerard, Maxime Girardin, Remi Grandemange, Jean-Claude Gregy, Joaquim Hatton, Thibaut Hingray, Thierry Hue, Arnaud Hurstel, Jean-Nöel Journot, Fabien Kilque, Lydie Lallement, Christian Lamboley, Manuel Lembke, Jean-Michel Letz, Vincent Lis, Olivier Marchand, Paul Massard, Yvan Mougel, Michel Munier, Louis-Michel Nageleisen, Yvan Nicolas, Christian Oberle, Pascal Perrotey-Doridant, Christian Philipps, François Rey-Demaneuf, Dorian Toussaint, Jean-Marie Triboulot, Bruno Vaxelaire, Laurent Verard, Jean-Lou Zimmermann, and Alice Zimmermann. We also thank Jacob Höglund and the other anonymous referee for their constructive comments that helped to improve the manuscript.

