## APPENDIX A for "Kin-dependent dispersal influences relatedness and genetic structuring in a lek system"

**APPENDIX A: study area, capture-recapture survey, and DNA analyses**

### 1. Study site, sampling design and sample collection

The Vosges Mountains are dominated by a south–north oriented ridge and small valleys separating low-altitude mountains. Until the 1970s, capercaillie distribution range extended to low altitude forests (400–500 m a.s.l.) dominated by Silver fir (*Abies alba*), Beech (*Fagus sylvatica*) and Scot pine (*Pinus sylvestris*), with a dense Bilberry (*Vaccinium myrtillus*) cover. Capercaillie distribution range contracted and the species nowadays persists at higher altitude (800–1250 m a.s.l.), in disconnected patches of mixed forest dominated by Silver fir, with Beech, Maple (*Acer* *sp.*) and Norway spruce (*Picea abies*). Capercaillie is also found at the subalpine range in Beech dominated forests and above tree line in moorlands dominated by ericaceous shrubs, where females find suitable habitat for the rearing of their broods. Lekking arenas are generally localised at the edges of moorlands dominated by ericaceous shrubs or moors. Scot pine needles is the preferred food source during winter, substituted by Silver fir needle in areas where Scot pine is absent (Lefranc & Preiss 2008).

As part of the routine monitoring of capercaillie, volunteers of the Groupe Tetras Vosges (hereafter agents) counted individuals in lekking arenas from lookouts. Once the birds had left the arenas, agents searched and collected faeces and feathers within a 400 m radius around lekking arenas. Prospections were repeated at one-month intervals between March and June. Agents also prospected newly established yet unstable lekking arenas and those historically occupied by the species.

Faeces were collected in 50 mL labelled screw cap tubes filled with 25 mL silica gel. Sampling date and coordinates of the sampling location were recorded. Tubes were stored at -20°C and kept frozen upon analysis. Faeces are considered waste products and are not covered by CITES (CITES Resolution Conf. 9.6, Rev CoP16). Nonetheless, importing animal by-products from France into Switzerland required an authorisation from the Swiss Federal Food Safety and Veterinary Office (Authorisation n° 1938/16).

### 2. DNA extraction

Strict laboratory procedures were adopted to control for potential sources of genotyping errors. Pre- and post-PCR experiments were conducted in separate rooms. Samples were manipulated with cleaned forceps (washed for 5 min in 10 % bleach and rinsed for 5 min in water) and, when required, cut using a sterile scalpel blade. Aerosol-resistant tips were used at all pipetting steps. We included 1–2 negative controls per batch of samples to control for cross-samples contamination or contamination of reagents. We extracted DNA from faecal samples following manufacturer recommendations, modified as described below. We used single-tubes to process < 22 samples and 96-plates to extract larger number of samples. DNA was eluted in 2 x 75 µL TE.

#### *Single-tube protocols (Qiagen Stool Mini Kit)*

Before manipulating the samples, we pipetted 2.7 ml of Buffer ASL (Qiagen) into a 15 ml tube, labelled and filled reaction tubes with reagents [Inhibitex tablet (Qiagen) and 20 µL proteinase K (Qiagen)], when required.

We cut 100 mg of faeces (up to 400 mg if the samples were moist) into pre-filled 15 mL tube, avoiding the white urea-rich part of the sample and incubated the samples overnight at room temperature. Tubes were thoroughly agitated (vortex) for 5–10 s to release the epithelial cells lining from the surface of the samples (do not disintegrate the sample as this releases inhibitors in the solution). We then transferred the supernatant into a pre-labelled 2 mL tube and centrifuge 1 min at 20000 g to pellet the particles.

#### *Single-tube protocols (Stratec PSP Spin Stool DNA Kit)*

Before manipulating the samples, we pipetted 2.0 ml of Lysis Buffer (Stratec) into a 15 ml tube, labelled and filled the reaction tubes with reagents 25 µL proteinase K (Stratec) when required.

We cut 100 mg of faeces (up to 400 mg if the samples were moist) into pre-filled 15 mL tube, avoiding the white urea-rich part of the sample. The tubes were shaken at 500 rpm for 2h to overnight at room temperature. We then transferred the supernatant into a pre-labelled 2 mL tube and centrifuge 1 min at 13400 g to pellet the particles.

#### *Plate protocol (Zymo ZR-96 Fecal DNA kit)*

Before manipulating the samples, we labelled and filled a 2 mL reaction tube with 200 µL Lysis Buffer I (50 mM Tris-HCl, 20 mM EDTA and 2 % w/v SDS). We added 40 mg of faecal sample (80–100 mg if the sample was moist) and grinded the sample with a metal spatula, added 200 µL Lysis Buffer D (Zymo) and grinded again the sample. Samples were incubated overnight (12–16 h) at room temperature. After brief centrifugation, the supernatant was pipetted into the extraction plate (including solid material increased DNA yield), with 100 µL each of Lysis Buffer I and D.

### 3. Individual identification

We amplified a fragment of the chromo-helicase gene for molecular sexing of the birds, using modified primers 1237 (Kahn et al. 1998) and P3 (Griffiths et al. 1998), and 19 microsatellite markers in two multiplexes (Table 1). Reactions were set in 10 µL volume containing 1x Type-it Multiplex kit (Qiagen), 1 µM MgCl_2_, 0.05 µL of each primer and 1–2 µL DNA template. Thermal cycling consisted of an activation step at 95°C for 5 min, 37 cycles of [94°C for 30 s, 54°C for 2 mn and 72°C for 30 s] and a final elongation step at 72°C for 15 min. DNA samples were amplified in four independent PCRs (multitube approach, Taberlet et al. 1996). We also amplified DNA from a known individual as a positive control in all PCR batches.

We mixed 1 µL of diluted PCR products (adding 20 µL ddH_2_O) with 7 µL (if loading a 384-well plates) or 10 µL (if loading a 96-well plates) HiDi Formamide (MCLab) and 2 % v/v internal size standard [GeneScan LIZ 500 (Thermo Fisher Scientific) or Orange Size Standard (MCLab)]. Fragment electrophoresis was conducted on ABI 3130 Genetic Analyzer (Thermo Fisher Scientific).

We used genemarker (SoftGenetics) to control for accurate scoring of internal size standard peaks. Samples showing low quality peaks were re-analysed. Laboratory conditions may affect electrophoretic mobility of PCR fragments (Davison & Chiba 2003) and induce genotyping errors. We used a reference individual of known genotype (positive PCR control) to ensure that allele sizing was consistent among runs. Alleles observed in ≥ 3 PCR replicates were coded as reliable and those observed twice were coded as low quality. Alleles observed once were ignored. Samples showing missing data at ≤ 4 microsatellite loci were amplified in 4–8 additional PCRs and we re-extracted samples showing missing data at > 5 loci. Sample genotypes showing missing data at more than four loci or low quality index (Miquel et al. 2006) after the second round of PCR amplification were excluded.

We grouped consensus genotypes and controlled mismatching alleles between sample genotypes mismatching at 1–3 loci. Reliable consensus genotypes were used to compute the rates of observed false alleles (FA) and allelic dropout (ADO) per loci. We estimated the probability that two random individuals share the same multi-locus genotype (*PI*) also accounting for the presence of siblings (*PI*_sibs_) using genalex (Peakall & Smouse 2012).

### 4. Genotyping success and sample size

We collected 1347 samples during the 6-years study, of which we excluded 112 (8.3 %) samples missing information on sampling date or location. Not all samples were amplified at the total set of 19 microsatellite loci and we therefore used a subset of 12 loci (ADL142, ADL184, ADL230, BG15, BG16, BG18, LEI098, TuT1, TuT2, TuT3, TuT4 and molecular sexing) shared among all samples. We could determine a reliable genotype in 962 (77.9 %) out of 1235 samples. Genotyping success ranged from 67.7 % (2013) to 86.3 % (2010; Table 2). We estimated rates of genotyping errors from mismatches between replicated genotypes in comparison to a consensus genotypes (Miquel et al. 2006). Rate of allelic dropout ranged from 0.14 (ADL142) to 0.23 (TuT4), and rate of false alleles and PCR artefacts were below 6 %.

We identified 132 individuals: 59 females, 70 males and three additional individuals for whom sex could not be determined. Numbers of individuals identified annually ranged from 64 (2010) to 33 (2014; Table 2). Most samples (75.9 %) were collected at eleven lek sites and during the breeding season, when 109 individuals were identified, including 61 males and 48 females. The remaining individuals were detected outside of the breeding season or in their wintering ranges and therefore excluded from CMR analyses. For molecular analyses, we analyzed the genetic data from 51 males and 41 females, keeping the individuals with no missing data (i.e. two alleles detected per individuals for all the markers).

Table A2. Number of samples collected per year between 2010 and 2015. 112 samples without information on sampling location or date were excluded. We also indicate annual numbers (and proportion) of samples successfully genotyped, and numbers of individuals (females/males) observed at lekking sites and during the breeding season

| Year | Collected | Genotyped | Selected |
| --- | --- | --- | --- |
| 2010 | 249 | 215 (86.3 %) | 69 (29/40) |
| 2011 | 178 | 129 (72.5 %) | 28 (10/18) |
| 2012 | 213 | 159 (74.6 %) | 48 (23/35) |
| 2013 | 201 | 136 (67.7 %) | 39 (14/25) |
| 2014 | 136 | 108 (79.4 %) | 30 (9/21) |
| 2015 | 258 | 215 (83.3 %) | 52 (17/35) |
| Total | **1347** | **962 (77.9 %)** | **117 (52/65)** |

Table A1. Nineteen microsatellites and a CHD-gene fragment were amplified, of which 12 loci (indicated by an asterisk) were used in the present study. We indicate locus name, multiplex number, fluorescent dye (Blue: FAM, Yellow: VIC^TM^/ATTO 532, Red: PET^TM^/ATTO 565, Green: NED^TM^/ATTO 550), number of alleles and allele range, forward and reverse sequences, GenBank accession number and reference.

| Locus | Mplex | #Allele (size-range) | Forward sequence (5’–3’) | Reverse sequence (5’–3’) | GeneBank #accession | Reference |
| --- | --- | --- | --- | --- | --- | --- |
| ADL184* | 1 | 2 (114–116) | Blue-GCCTCCTCACCCACAAAACC | TCAGTAACACCACGAATGCC | G01606 | Cheng unpublished |
| ADL230* | 1 | 2 (109–111) | Red-GCCAAATAGTAATCCACTGC | TCGCTCTTGCCATTGTAAGT | G01650 | Cheng unpublished |
| BG15* | 1 | 3 (135–143) | Yellow-AAATATGTTTGCTAGGGCTTAC | TACATTTTTCATTGTGGACTTC | AF381549 | Piertney & Höglund (2001) |
| BG16* | 1 | 4 (166–178) | Red-GTCATTAGTGCTGTCTGTCTATCT | TGCTAGGTAGGGTAAAAATGG | AF381550 | Piertney & Höglund (2001) |
| BG18* | 1 | 6 (183–207) | Yellow-CCATAACTTAACTTGCACTTTC | CTGATACAAAGATGCCTACAA | AF381551 | Piertney & Höglund (2001) |
| LEI098* | 1 | 5 (142–158) | Blue-CAGTTAGCAGAGATTTTCCTAC | TGCCACTGATGCTGTCACTG | X82860 | Gibbs et al 1997 |
| TuT1* | 1 | 4 (199–219) | Blue-GGTCTACATTTGGCTCTGACC | ATATGGCATCCCAGCTATGG | AF254653 | Segelbacher et al (2000) |
| TuT2* | 1 | 2 (157–161) | Yellow-CCGTGTCAAGTTCTCCAAAC | TTCAAAGCTGTGTTTCATTAGTTG | AF254654 | Segelbacher et al (2000) |
| TuT3* | 1 | 3 (151–159) | Green-CAGGAGGCCTCAACTAATCACC | CGATGCTGGACAGAAGTGAC | AF254655 | Segelbacher et al (2000) |
| TuT4* | 1 | 3 (171–187) | Green-GAGCATCTCCCAGAGTCAGC | TGTGAACCAGCAATCTGAGC | AF254656 | Segelbacher et al (2000) |
| 1237rc^1^/P3rc^2^* | 1+2 | Z (244), W (269) | Blue- RATGAGAAACTGTGCAAAACAG | GGARTCACTATCAGATCCAGAATATC |  |  |
| ADL142* | 2 | 3 (211–217) | Yellow-CAGCCAATAGGGATAAAAGC | CTGTAGATGCCAAGGAGTGC | G01567 | Cheng unpublished |
| BG10 | 2 | 3 (190–202) | Green-ATGTTTCATGTCTTCTGGAATAG | ATTTGGTTAGTAACGCATAAGC | AF381546 | Piertney & Höglund (2001) |
| BG12 | 2 | 4 (150–178) | Red-TCTCCTTCTAAACCAGTCATTC | TAGTTTCCACAGAGCACATTG | AF381547 | Piertney & Höglund (2001) |
| BG20 | 2 | 3 (123­131) | Green-AAGCACTTACAATGGTGAGGAC | TATGTTTTCCTTTTCAGTGGTATG | AF381553 | Piertney & Höglund (2001) |
| BG6 | 2 | 2 (194–198) | Blue-AAAGAGGCAAGCACTCACAATG | CCCTTGGAATATCCTTTAACAAAAC |  | Piertney & Höglund (2001) |
| sTuD1 | 2 | 5 (155–174) | Blue-ATTTGCCAGGAAACTTGCTC | CCTTTGCCTCCTTATGAAATCC | AF254644 | Jacob et al (2009)^3^ |
| sTuD3 | 2 | 3 (82–92) | Blue-CAAGGGGAAAATATGTGTGTG | TGTCAAGATATTTCAAGCCTTTG | AF254646 | Jacob et al (2009)^3^ |
| sTuD6 | 2 | 3 (168–180) | Yellow-AGCCTTTTACTGCACTACTTGC | GGTGTGTGGGAAATGAGGAC | AF254649 | Jacob et al (2009)^3^ |
| sTuD7 | 2 | 4 (91–105) | Yellow-GGGTCATTAGGCAGAGCTTTC | CCTGCATCATTCCAAATGTC | AF254650 | Jacob et al (2009)^3^ |

**^1^ modified from Kahn et al.** (Kahn et al. 1998)

**^2^ modified from Griffiths et al.** (Griffiths et al. 1998)

^3^ **modified from Segelbacher et al.** (Segelbacher et al. 2000)
