## APPENDIX B for "Kin-dependent dispersal influences relatedness and genetic structuring in a lek system"

**APPENDIX B: States and events of Multievent Capture-Recapture models**

Table B1. Quantifying the proportion of dispersers per generation: model states and events

| Notation | State or event description |
| --- | --- |
| States |  |
| So | Individual with a disperser strategy, stayed in the same lek at *t* that the one occupied at *t*–1, and not captured at *t* |
| oS+ | Individual with a disperser strategy, not captured at *t*–1, stayed in the same lek at *t* that the one occupied at *t*–1, and captured at *t* |
| +S+ | Individual with a disperser strategy, captured at *t*–1, stayed in the same lek at *t* that the one occupied at *t*–1, and captured at *t* |
| Mo | Individual with a disperser strategy, moved to another lek between *t*–1 and *t*, and not captured at *t* |
| oM+ | Individual with a disperser strategy, not captured at *t*–1, moved to another lek between *t*–1 and *t*, and captured at *t* |
| +M+ | Individual with a disperser strategy, captured at *t*–1, moved to another lek between *t*–1 and *t*, and captured at *t* |
| Ro | Individual with a fully resident strategy, stayed in the same lek at *t* that the one occupied at *t*–1, and not captured at *t* |
| oR+ | Individual with a fully resident strategy, not captured at *t*–1, stayed in the same lek at *t* that the one occupied at *t*–1, and captured at *t* |
| +R+ | Individual with a fully resident strategy, captured at *t*–1, stayed in the same lek at *t* that the one occupied at *t*–1, and captured at *t* |
| D | Dead |
| Events |  |
| 0 | Not captured at t |
| 1 | Captured at *t*, not captured at *t*–1 |
| 2 | Captured at *t*, in the same lek at *t* that the one occupied at *t*–1 |
| 3 | Captured at *t*, moved to another lek between *t*–1 and *t* |

Table B2. Quantifying annual dispersal rates and distances: model states and events.

| Notation | State or event description |
| --- | --- |
| So | Not captured at *t*, stayed in the same lek at *t* that the one occupied at *t*–1 |
| oS+ | Not captured at *t*–1, captured at *t*, and stayed in the same lek at *t* that the one occupied at *t*–1. |
| +S+ | Captured at *t*–1 and *t*, stayed in the same lek at *t* that the one occupied at *t*–1, and captured at *t* |
| M1o | Not captured at *t*, moved to another lek between *t*–1 and *t*, and recipient lek located within the distance class 1 |
| M1+ | Not captured at *t*–1, captured at *t*, moved to another lek between *t*–1 and *t*, and recipient lek located within the distance class 1 |
| +M1+ | Captured at *t*–1 and *t*, moved to another lek between *t*–1 and *t*, and recipient lek located within the distance class 1 |
| M2o | Not captured at *t*, moved to another lek between *t* – 1 and *t*, and recipient lek located within the distance class 2 |
| M2+ | Not captured at *t*–1, captured at *t*, moved to another lek between *t* – 1 and *t*, and recipient lek located within the distance class 2 |
| +M2+ | Captured at *t*–1 and *t*, moved to another lek between *t*–1 and *t*, and recipient lek located within the distance class 2 |
| M3o | Not captured at *t*, moved to another lek between *t* – 1 and *t*, and recipient lek located within the distance class 2 |
| M3+ | Not captured at *t*–1, captured at *t*, moved to another lek between *t* – 1 and *t*, and recipient lek located within the distance class 2 |
| +M3+ | Captured at *t*–1 and *t*, moved to another lek between *t*–1 and *t*, and recipient lek located within the distance class 2 |
| M4o | Not captured at *t*, moved to another lek between *t* – 1 and *t*, and recipient lek located within the distance class 4 |
| M4+ | Not captured at *t*–1, captured at *t*, moved to another lek between *t* – 1 and *t*, and recipient lek located within the distance class 4 |
| +M4+ | Captured at *t*–1 and *t*, moved to another lek between *t*–1 and *t*, and recipient lek located within the distance class 4 |
| M5o | Not captured at *t*, moved to another lek between *t* – 1 and *t*, and recipient lek located within the distance class 4 |
| M5+ | Not captured at *t*–1, captured at *t*, moved to another lek between *t* – 1 and *t*, and recipient lek located within the distance class 4 |
| +M5+ | Captured at *t*–1 and *t*, moved to another lek between *t*–1 and *t*, and recipient lek located within the distance class 4 |
| D | Dead |
| Events |  |
| 0 | Not captured at t |
| 1 | Captured at *t*, in the same lek at *t* that the one occupied at *t*–1 |
| 2 | Captured at *t*, not captured at *t*–1 |
| 3 | Captured at *t*, moved to another lek between *t*–1 and *t*, recipient lek located within the distance class 1 |
| 4 | Captured at *t*, moved to another lek between *t*–1 and *t*, recipient lek located within the distance class 2 |
| 5 | Captured at *t*, moved to another lek between *t*–1 and *t*, recipient lek located within the distance class 3 |
| 6 | Captured at *t*, moved to another lek between *t*–1 and *t*, recipient lek located within the distance class 4 |
| 7 | Captured at *t*, moved to another lek between *t*–1 and *t*, recipient lek located within the distance class 5 |
