## APPENDIX C for "Kin-dependent dispersal influences relatedness and genetic structuring in a lek system"

**APPENDIX C: model selection procedure for capture-recapture analyses**

Table C1. Quantifying non-effective dispersal rate per generation: model selection procedure. Model parameters may vary between sexes and years. μ = proportion of dispersers (i.e. individuals that dispersed at least one time during their lifetime) per generation, ϕ = survival probability, ψ = dispersal probability, *p* = recapture probability, k = number of model parameters, Deviance = residual deviance, AICc = Akaike information criterion adjusted for small sample size, w = AICc weight.

| Model | k | Deviance | AICc | w |
| --- | --- | --- | --- | --- |
| μ(.), ϕ(.), ψ(.), *p*(SEX) | 5 | 525.25 | 535.52 | 0.22 |
| μ(.), ϕ(SEX), ψ(.), *p*(SEX) | 6 | 523.56 | 535.92 | 0.18 |
| μ(SEX), ϕ(.), ψ(.), *p*(SEX) | 6 | 524.07 | 536.44 | 0.14 |
| μ(SEX), ϕ(SEX), ψ(.), *p*(SEX) | 7 | 522.37 | 536.86 | 0.11 |
| μ(.), ϕ(.), ψ(SEX), *p*(SEX) | 6 | 525.20 | 537.57 | 0.08 |
| μ(SEX), ϕ(.), ψ(SEX), *p*(SEX) | 7 | 523.45 | 537.94 | 0.06 |
| μ(.), ϕ(SEX), ψ(SEX), *p*(SEX) | 7 | 523.50 | 537.99 | 0.06 |
| μ(SEX), ϕ(SEX), ψ(SEX), *p*(SEX) | 8 | 521.75 | 538.38 | 0.05 |
| μ(.), ϕ(.), ψ(.), *p*(YEAR+SEX) | 9 | 521.18 | 539.98 | 0.02 |
| μ(SEX), ϕ(.), ψ(.), *p*(YEAR+SEX) | 10 | 520.00 | 540.98 | 0.01 |
| μ(.), ϕ(SEX), ψ(.), *p*(YEAR+SEX) | 10 | 520.09 | 541.07 | 0.01 |
| μ(SEX), ϕ(SEX), ψ(.), *p*(YEAR+SEX) | 11 | 518.91 | 542.09 | 0.01 |
| μ(.), ϕ(.), ψ(SEX), *p*(YEAR+SEX) | 10 | 521.13 | 542.11 | 0.01 |
| μ(SEX), ϕ(.), ψ(SEX), *p*(YEAR+SEX) | 11 | 519.38 | 542.56 | 0.01 |
| μ(.), ϕ(SEX), ψ(SEX), *p*(YEAR+SEX) | 11 | 520.04 | 543.22 | 0.00 |
| μ(SEX), ϕ(SEX), ψ(SEX), *p*(YEAR+SEX) | 12 | 518.29 | 543.69 | 0.00 |
| μ(.), ϕ(.), ψ(.), *p*(.) | 4 | 534.96 | 543.13 | 0.00 |
| μ(SEX), ϕ(.), ψ(.), *p*(.) | 5 | 533.77 | 544.03 | 0.00 |
| μ(.), ϕ(.), ψ(SEX), *p*(.) | 5 | 534.90 | 545.16 | 0.00 |
| μ(.), ϕ(SEX), ψ(.), *p*(.) | 5 | 534.95 | 545.21 | 0.00 |
| μ(SEX), ϕ(.), ψ(SEX), *p*(.) | 6 | 533.15 | 545.52 | 0.00 |
| μ(SEX), ϕ(SEX), ψ(.), *p*(.) | 6 | 533.76 | 546.13 | 0.00 |
| μ(.), ϕ(SEX), ψ(SEX), *p*(.) | 6 | 534.89 | 547.26 | 0.00 |
| μ(.), ϕ(.), ψ(.), *p*(YEAR) | 8 | 530.99 | 547.62 | 0.00 |
| μ(SEX), ϕ(SEX), ψ(SEX), *p*(.) | 7 | 533.14 | 547.63 | 0.00 |
| μ(SEX), ϕ(.), ψ(.), *p*(YEAR) | 9 | 529.80 | 548.60 | 0.00 |
| μ(.), ϕ(.), ψ(SEX), *p*(YEAR) | 9 | 530.93 | 549.73 | 0.00 |
| μ(.), ϕ(SEX), ψ(.), *p*(YEAR) | 9 | 530.97 | 549.76 | 0.00 |
| μ(SEX), ϕ(.), ψ(SEX), *p*(YEAR) | 10 | 529.18 | 550.16 | 0.00 |
| μ(SEX), ϕ(SEX), ψ(.), *p*(YEAR) | 10 | 529.78 | 550.76 | 0.00 |
| μ(.), ϕ(SEX), ψ(SEX), *p*(YEAR) | 10 | 530.91 | 551.89 | 0.00 |
| μ(SEX), ϕ(SEX), ψ(SEX), *p*(YEAR) | 11 | 529.16 | 552.34 | 0.00 |

Table C2. Quantifying non-effective annual dispersal rates and distances: model selection procedure. Model parameters may vary between sexes and years. ϕ = survival probability, τ = departure probability, α = arrival probability, *p* = recapture probability, k = number of model parameters, Deviance = residual deviance, AICc = Akaike information criterion adjusted for small sample size, w = AICc weight.

| Model | k | Deviance | AICc | w |
| --- | --- | --- | --- | --- |
| φ(.), τ(SEX), α(.), *p*(SEX) | 9 | 583.97 | 602.76 | 0.16 |
| φ(.), τ(.), α(.), *p*(SEX) | 8 | 586.17 | 602.81 | 0.16 |
| φ(SEX), τ(SEX), α(.), *p*(SEX) | 10 | 582.27 | 603.25 | 0.12 |
| φ(SEX), τ(.), α(.), *p*(SEX) | 9 | 584.48 | 603.27 | 0.12 |
| φ(.), τ(SEX), α(SEX), *p*(SEX) | 10 | 582.74 | 603.72 | 0.10 |
| φ(.), τ(.), α(SEX), *p*(SEX) | 9 | 584.95 | 603.75 | 0.10 |
| φ(SEX), τ(SEX), α(SEX), *p*(SEX) | 11 | 581.05 | 604.22 | 0.08 |
| φ(SEX), τ(.), α(SEX), *p*(SEX) | 10 | 583.25 | 604.23 | 0.08 |
| φ(.), τ(SEX), α(.), *p*(YEAR+SEX) | 13 | 579.89 | 607.53 | 0.01 |
| φ(.), τ(.), α(.), *p*(YEAR+SEX) | 12 | 582.10 | 607.50 | 0.01 |
| φ(.), τ(SEX), α(SEX), *p*(YEAR+SEX) | 14 | 578.67 | 608.57 | 0.01 |
| φ(SEX), τ(SEX), α(.), *p*(YEAR+SEX) | 14 | 578.81 | 608.71 | 0.01 |
| φ(.), τ(.), α(SEX), *p*(YEAR+SEX) | 13 | 580.88 | 608.52 | 0.01 |
| φ(SEX), τ(.), α(.), *p*(YEAR+SEX) | 13 | 581.01 | 608.65 | 0.01 |
| φ(SEX), τ(SEX), α(SEX), *p*(YEAR+SEX) | 15 | 577.58 | 609.76 | 0.00 |
| φ(SEX), τ(.), α(SEX), *p*(YEAR+SEX) | 14 | 579.79 | 609.69 | 0.00 |
| φ(.), τ(SEX), α(.), *p*(.) | 8 | 593.67 | 610.30 | 0.00 |
| φ(.), τ(.), α(.), *p*(.) | 7 | 595.88 | 610.37 | 0.00 |
| φ(.), τ(SEX), α(SEX), *p*(.) | 9 | 592.44 | 611.24 | 0.00 |
| φ(.), τ(.), α(SEX), *p*(.) | 8 | 594.65 | 611.29 | 0.00 |
| φ(SEX), τ(SEX), α(.), *p*(.) | 9 | 593.66 | 612.46 | 0.00 |
| φ(SEX), τ(.), α(.), *p*(.) | 8 | 595.87 | 612.50 | 0.00 |
| φ(SEX), τ(SEX), α(SEX), *p*(.) | 10 | 592.44 | 613.41 | 0.00 |
| φ(SEX), τ(.), α(SEX), *p*(.) | 9 | 594.64 | 613.44 | 0.00 |
| φ(.), τ(SEX), α(.), *p*(YEAR) | 12 | 589.70 | 615.10 | 0.00 |
| φ(.), τ(.), α(.), *p*(YEAR) | 11 | 591.91 | 615.09 | 0.00 |
| φ(.), τ(SEX), α(SEX), *p*(YEAR) | 13 | 588.48 | 616.12 | 0.00 |
| φ(.), τ(.), α(SEX), *p*(YEAR) | 12 | 590.69 | 616.08 | 0.00 |
| φ(SEX), τ(SEX), α(.), *p*(YEAR) | 13 | 589.68 | 617.32 | 0.00 |
| φ(SEX), τ(.), α(.), *p*(YEAR) | 12 | 591.89 | 617.29 | 0.00 |
| φ(SEX), τ(SEX), α(SEX), *p*(YEAR) | 14 | 588.46 | 618.36 | 0.00 |
| φ(SEX), τ(.), α(SEX), *p*(YEAR) | 13 | 590.66 | 618.30 | 0.00 |

**Yearly-specific dispersal rates in the population of Western capercaillie (*Tetrao urogallus*).**

We examined how departure rate fluctuated between years by comparing the AICc of the two following models, (φ(.), τ(.), α(.), *p*(SEX)) and (φ(.), τ(YEAR), α(.), *p*(SEX)). In the first model, departure rate was hold constant (τ(.)) whereas it varied between years in the second model (τ(YEAR)). The model in which departure was hold constant was better supported by the data (AICc = 602.81) than the model with a year effect (AICc = 605.48). This result indicates lowly variable departure rates between years (see Figure C1).

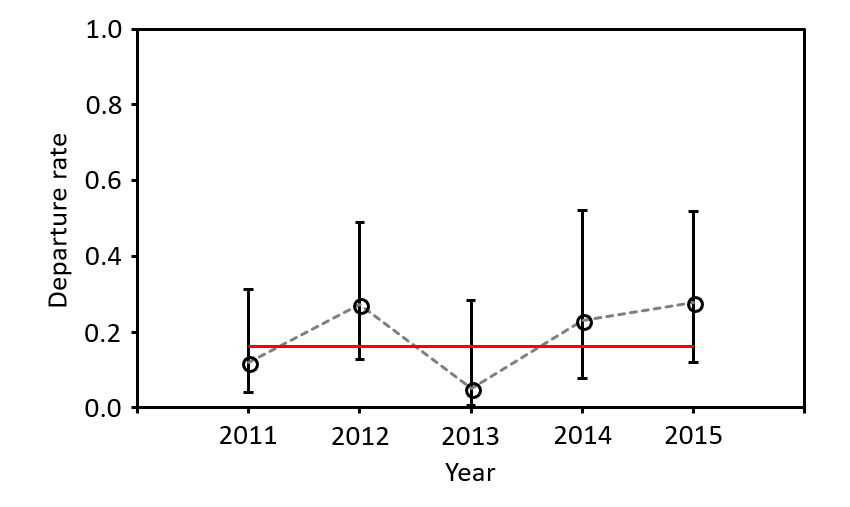

Figure C1. Yearly-specific dispersal rates in the population of Western capercaillie (*Tetrao urogallus*). We show yearly-specific departure rates and their 95% CI extracted from the model (φ(.), τ(YEAR), α(.), *p*(SEX)). The red line corresponds to constant departure rate extracted from the model (φ(.), τ(.), α(.), *p*(SEX)).
