## APPENDIX D for "Kin-dependent dispersal influences relatedness and genetic structuring in a lek system"

**APPENDIX D: Population genetic structure**

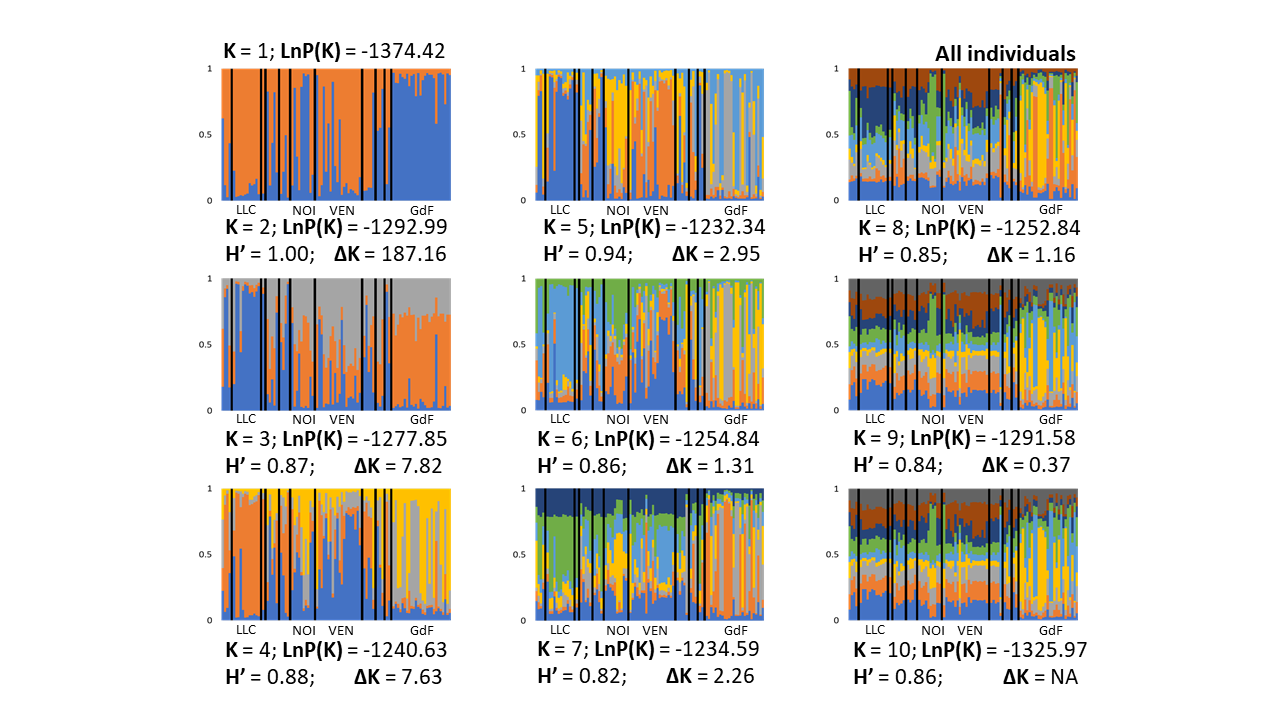

Fig. D1. STRUCTURE outputs for all the individuals (males and females).

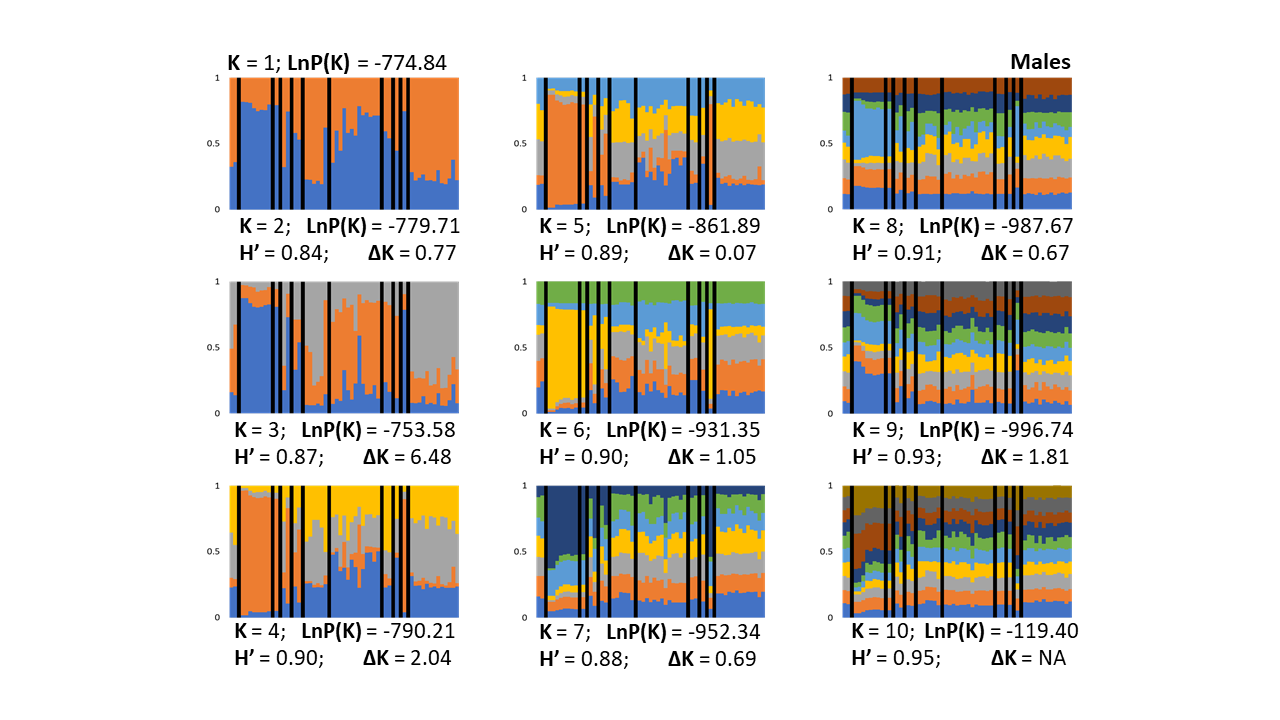

Fig. D2. STRUCTURE outputs for males only.

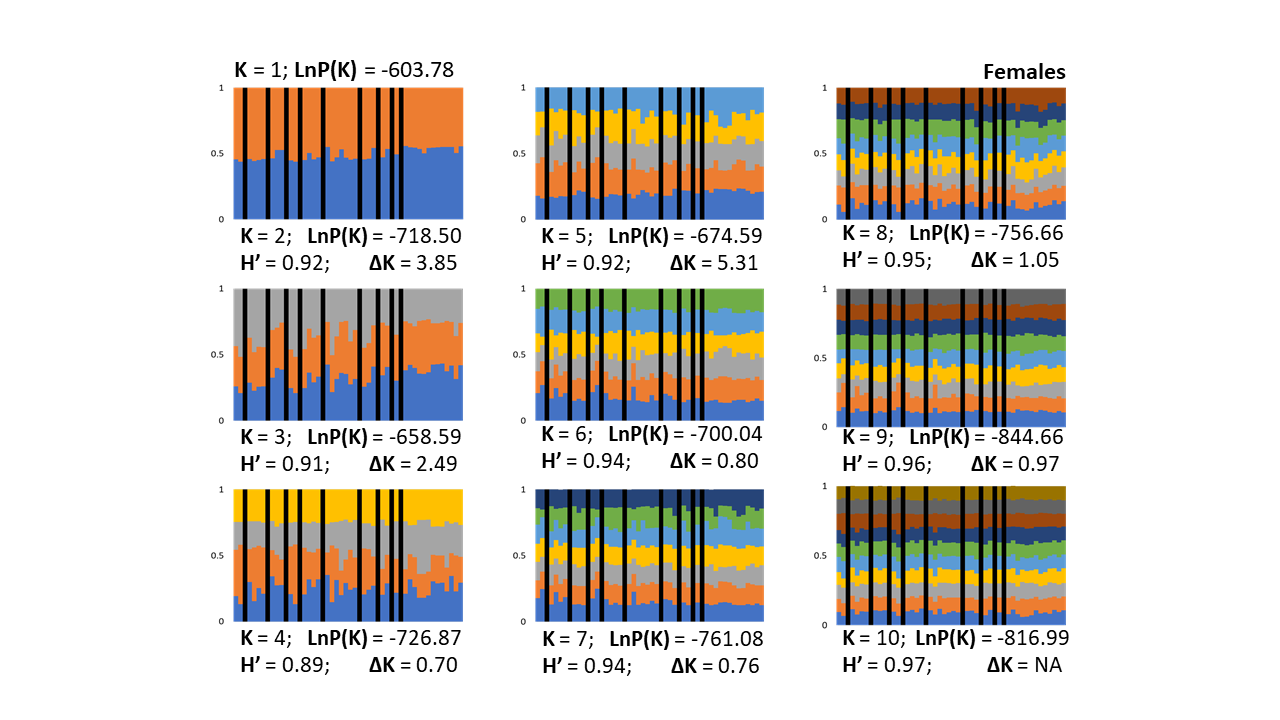

Fig. D3. STRUCTURE outputs for females only.

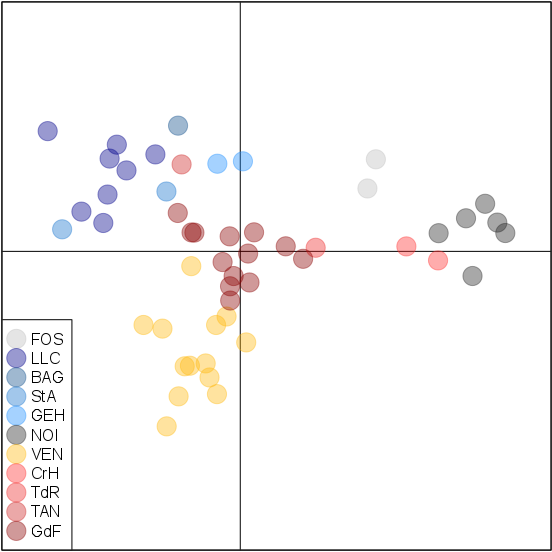

Fig. D4. DAPC for males only.

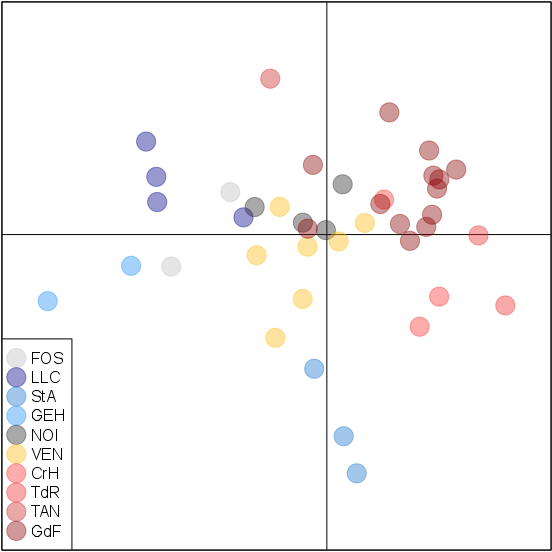

Fig. D5. DAPC for females only.

Table D1. Results of the simulations performed in CO-ANCESTRY. Our analyses revealed that the estimator LynchRd (shown in grey) was the most accurate for our dataset.

|  | **TrioML** | **Wang** | **LynchLi** | **LynchRd** | **Ritland** | **QuellerGt** | **DyadML** | **True Value** |
| --- | --- | --- | --- | --- | --- | --- | --- | --- |
| **Mean** | 0,26 | 0,22 | 0,22 | 0,22 | 0,21 | 0,21 | 0,3 | 0,23 |
| **Variance** | 0,05 | 0,12 | 0,12 | 0,1 | 0,15 | 0,11 | 0,06 | 0,04 |
| **MSE** | 0,04 | 0,08 | 0,08 | 0,06 | 0,12 | 0,07 | 0,04 | NA |
| **r Pearson** | 0,6 | 0,57 | 0,57 | 0,61 | 0,45 | 0,57 | 0,62 | NA |
