## APPENDIX E for "Kin-dependent dispersal influences relatedness and genetic structuring in a lek system"

**APPENDIX E: Influence of landscape predictors on genetic variation**

**A. Details on landscape predictors.**

*Dtopo* was computed as the cumulated length of segments that would be travelled by an individual flying in straight line from a point A to a point B while following topographic relief. From a Digital Elevation Model (DEM) at a 25m resolution, we computed the effective travelled distance *d* across each pixel *i* crossed by an individual along its straight line trajectory using the Pythagorean Theorem as follows: $d_{i}= \sqrt{\left( x_{i-1}-x_{i} \right)^{2}+\left( y_{i-1}-y_{i} \right)^{2}}$ , with $\left( x_{i-1}-x_{i} \right)=25$ (the length of a pixel), $y_{i-1}$ the altitude at the previous pixel along the trajectory and $y_{i}$ the altitude at the focal pixel. Dtopo was then computed as the sum of distances $d_{i}$ along the trajectory.

To compute *Dslope*, we first created a slope raster map at a 25m resolution from the DEM using the Spatial Analyst toolbox in ArcGis 10.2. The layer was rescaled to range from 1 to 100 and we used CIRCUITSCAPE 3.5.8 to compute pairwise effective distances between individual locations, with the raster coded in resistances (higher slope values denoting greater resistance to movement).

To compute *Dridge*, we first applied the Focal Statistics tool from the Spatial Analyst toolbox to compute, for each pixel from the DEM, the mean elevation values within a 1000m neighborhood. We then subtracted the Focal Stats layer from the original DEM and identified ridges as pixels associated with a positive value (pixels above the average elevation of their neighborhood). Altitude values of ridge pixels were rescaled to range from 1(highest elevation) to 100 (lowest elevation) whereas non-ridge pixels were systematically set to 100. We finally used CIRCUITSCAPE 3.5.8 to compute pairwise effective distances between individual locations, with the raster coded in resistances (non-ridge areas denoting greater resistance to movement).

**B. Identification of the main contributors to the variance in pairwise measures of genetic differentiation.**

We coupled multiple linear regressions on standardized data at the optimal scale of analysis (see main text for details) with commonality analyses (CA; Ray-Murkherjee et al. 2014) to identify the main contributors to the variance in the dependent variables after sequential removing of suppressors and unnecessary predictors. Predictors were identified as suppressors when their unique contribution was (almost) totally counterbalanced by a negative commonality coefficient (classical and reciprocal suppression) or when standardized regression coefficients and structure coefficients were of opposite signs (cross-over suppression; Paulhus et al. 2004, Prunier et al. 2017). Predictors were identified as unnecessary when their unique contribution (U) was null (or when the lower bound of 95 % confidence intervals CI around U was null), indicating that they only contributed to the variance in the dependent variable because of their synergistic association with one or several other predictors (Prunier et al., 2015). The 95 % CI around beta coefficients β and unique contributions U were computed from bootstrap resampling (1000 iterations).

For each sex (M and F), the following table provides details about runs of identification of unnecessary predictors (in synergistic association with other predictors) and suppressors in full models (see main text for details). Are provided: typical results of the different runs of multiple linear regressions (model fit R², predictors Pred, structure coefficients rs and standardized coefficents β), along with additional parameters derived from CA: unique, common and total contributions of predictors to the variance in dependent variable (U, C and T), as well as 95% confidence intervals about U (UCI_low_ and UCI_up_) as computed from bootstrap (1000 iterations). The rationale for withdrawal of predictors (Ra) is the following: S: synergistic association with other predictors (UCI_low_ = 0); CO: Cross-over suppression. In bold: parameters allowing the identification of unnecessary predictors and suppressors.

| Sex | Run | R² | Pred | rs | β | Unique | UCI_low_ | UCI_up_ | Common | Total | %U | Ra |
| --- | --- | --- | --- | --- | --- | --- | --- | --- | --- | --- | --- | --- |
| M | 1 | 0.308 | *ED* | 0.932 | 0.242 | 0.007 | **0.000** | 0.024 | 0.261 | 0.267 | 0.022 | S |
|  |  |  | *Dtopo* | 0.916 | 0.246 | 0.014 | 0.002 | 0.038 | 0.244 | 0.258 | 0.046 |  |
|  |  |  | *Dridge* | **0.819** | **-0.096** | 0.002 | 0.000 | 0.016 | 0.205 | 0.207 | 0.006 | CO |
|  |  |  | *Dslope* | 0.817 | 0.223 | 0.027 | 0.009 | 0.051 | 0.179 | 0.205 | 0.087 |  |
|  | 2 | 0.301 | *Dtopo* | 0.926 | 0.371 | 0.095 | 0.057 | 0.144 | 0.163 | 0.258 | 0.317 |  |
|  |  |  | *Dslope* | 0.827 | 0.248 | 0.043 | 0.019 | 0.073 | 0.163 | 0.205 | 0.142 |  |
| F | 1 | 0.054 | *ED* | 0.666 | 0.243 | 0.008 | **0.000** | 0.027 | 0.016 | 0.024 | 0.150 | S |
|  |  |  | *Dtopo* | 0.752 | 0.212 | 0.010 | 0.001 | 0.029 | 0.020 | 0.031 | 0.189 |  |
|  |  |  | *Dridge* | **0.294** | **-0.336** | 0.022 | 0.005 | 0.048 | -0.017 | 0.005 | 0.398 | CO |
|  |  |  | *Dslope* | 0.306 | 0.030 | 0.000 | **0.000** | 0.010 | 0.005 | 0.005 | 0.006 | S |
|  | 2 | 0.031 | *Dtopo* | 0.175 | 0.175 | / | / | / | / | / | / |  |
