## APPENDIX F for "Kin-dependent dispersal influences relatedness and genetic structuring in a lek system"

**APPENDIX F: Relatedness effect on first capture and departure**

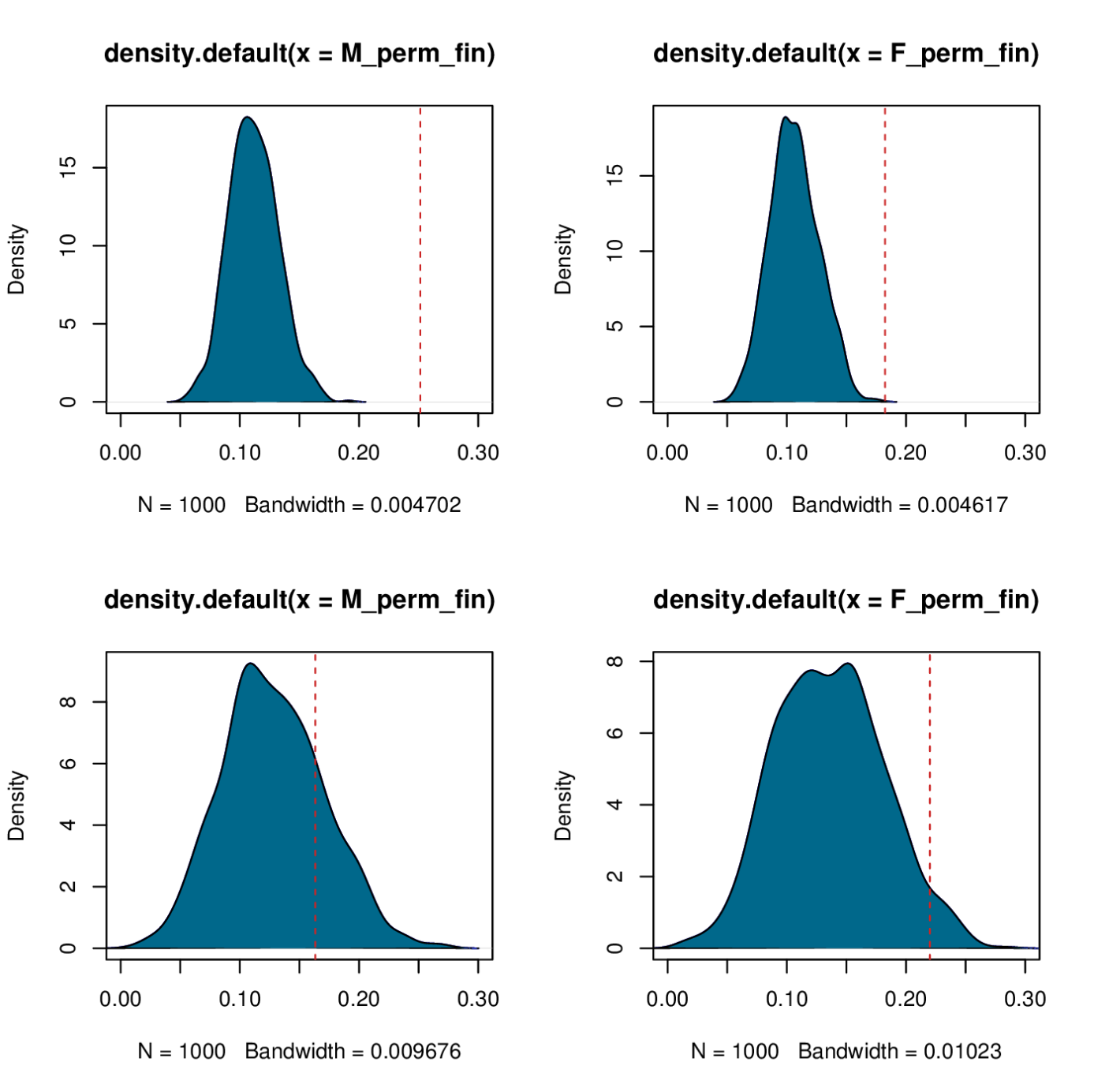

Fig. F1. Distribution of the mean of average relatedness within lek. Left: males only; Right: females only; Top: permutated first capture; Bottom: permutated departure. Red dash line represent original dataset value.
